# Longitudinal Neurocognitive Trajectories in a Large Cohort of Youth Who Use Cannabis: Combining Self-Report and Toxicology

**DOI:** 10.64898/2025.12.20.695698

**Authors:** Natasha E. Wade, Ryan M. Sullivan, Alexander L. Wallace, Rachel Visontay, Veronica Szpak, Krista M. Lisdahl, Marilyn A. Huestis, Priscila Dib Gonçalves, Hollie Byrne, Louise Mewton, Joanna Jacobus, Susan F. Tapert

## Abstract

Adolescents experience extensive neurocognitive development, with cannabis use potentially impacting developmental trajectories. Here, we comprehensively assess the influence of adolescent cannabis use onset on neurocognitive trajectories and consider how recent delta-9-tetrahydrocannabinol (THC) and cannabidiol (CBD) may influence neurocognition. We use the large, diverse longitudinal Adolescent Brain Cognitive Development (ABCD) Study dataset, combining self-reported substance use with objective toxicological tests (hair, urine, breath, oral fluid). Longitudinal mixed methods of the full cohort (n=11,036, ages 9-17; 47% Female/53% Male) investigate time-varying cannabis onset on neurocognitive performance. Primary model covariates include sociodemographics, family history of substance use disorder, prenatal substance exposure, early psychopathology, other substance use, and nesting for participant ID, study site, and family ID. Secondarily, in participants with repeat toxicological hair testing (n=645; 38% Female/62% Male) at ages 12-16, we consider the influence of THC v. CBD v.

Controls. Primary models included false discovery rate corrections (FDR-p<.05) while secondary models were interpreted at p<.01. Cannabis group interacted with age to show altered neurocognitive trajectories across domains (immediate recall and delayed memory, processing speed, inhibitory control, visuospatial processing, language, and working memory; βs=-0.11- - 0.52). Secondary models indicated hair-identified THC exposure*age predicted worse episodic memory than in Controls (β=-0.60, *p*=.007), with no difference between CBD exposed and Controls. Data suggest those who use cannabis show likely pre-existing better cognitive performance during late childhood, with reduced improvement or flattened trajectories over time. These neurocognitive trajectories in youth (ages 9-17) who initiate cannabis use were demonstrated after accounting for within-person change and numerous known confounds and improving accuracy in identifying cannabis use through incorporating toxicological measures. Continued monitoring of this cohort will clarify cannabinoid-cognition relationships into young adulthood, including the impact of timing of cannabis use initiation.

## Introduction

Over the past several decades, high rates of cannabis use in individuals ages 13-24 remained relatively stable^1^, but the legal status of cannabis changed in many US states, the potency of the primary psychoactive cannabinoid delta-9-tetrahydrocannabinol (THC) increased^2^, vaped inhalation increased, and public perception of risk decreased^3^. Critically, adolescents experience extensive brain and cognitive development^4–6^, making adolescents particularly vulnerable to the influence of cannabis^7^. This includes within the endocannabinoid system, a neuromodulatory system critical to brain development (e.g., neural progenitor proliferation, neurogenesis)^7,8^ and in which THC exerts its psychoactive effects. THC further influences dopaminergic, GABA, and glutamate activity across cortico-limbic regions, all which are key to healthy neurodevelopment during the neuroplastic period of adolescence^9^. Early longitudinal studies suggest neurocognitive deficits in select domains are associated with adolescent cannabis onset and escalation, such as poorer verbal learning and memory^10–16^, visuospatial functioning^11,12,14,16^, working memory^11,12,16,17^, inhibition and attention^12,15,16,18^, and psychomotor and processing speed^10,12,14,15^. However, these studies often do not fully account for important pre-substance use confounds (e.g., other substance use, family history, family factors, sociodemographics), include small sample sizes, or do not clearly identify the onset of cannabis use^17,19,20^, and not all identify relative cognitive decrements^21^.

Studies of adolescent cannabis use can also be limited by the accuracy of self-report, the primary method of identifying cannabis use in most studies. Recent studies comparing objective, toxicological measures with self-report in adolescents and young adults underestimated substance use prevalence by up to 60%^22^, or mis-identified up to 9% of participants as “controls” when they actually used substances^23^. Therefore, research on the sensitive topic of substance use in teens may at times present with bias in self-report, which impedes identifying substance use consequences. On balance, toxicological testing may not identify low-level (e.g., one or several uses) or use outside of range of detection (e.g., >3 months for hair testing), thus necessitating querying self-reported substance use. Incorporating toxicological measures, including hair cannabinoids, with self-report may improve accuracy in characterizing cannabis-neurocognition relationships^24,25^.

Accurate assessment of cannabis use histories is further complicated by the complexity of the plant that contains over 120 cannabinoids^26^, with the most consistent evidence demonstrating repeated THC exposure is linked with changes in CB1 density and downstream cognitive functioning^27^. Adult evidence demonstrates that cannabidiol (CBD), a CB1 cannabinoid receptor allosteric modulator, may offer mitigating influence on cognitive performance^28,29^ especially with chronic exposure^30^, though this is not always found^31,32^ and the impact on adolescents is unclear. Objective evidence of cannabinoid analyte concentration exposures may be more informative than relying on self-reported cannabis use episodes or potency, especially given individual differences in processing of cannabis due to genetics, rates of exposure, and other biological mechanisms^33–36^, in addition to inconsistencies in self-reported use^23^. The literature on specific cannabinoid constituents as related to cognitive functioning in the developing brain is exceptionally limited^37,38^, yet different cannabinoids may moderate clinical outcomes, as shown in studies considering psychopathology using hair samples in adolescents and young adults^25,39–41^. Thus, combining toxicological cannabinoid information with self-reported use may more accurately and thoroughly assess the impact of cannabis use onset on cognition and further investigate the influence of specific cannabinoid constituents (i.e., THC, CBD, the most studied cannabinoids) as potential explanatory mechanisms.

Here, we comprehensively assess the influence of adolescent cannabis use onset on neurocognitive trajectories. We probe potential mechanisms by investigating specific cannabinoid content through toxicological hair testing to examine whether use of certain cannabinoids may predict differential trajectories. Leveraging data from the Adolescent Brain Cognitive Development (ABCD) Study–a large and richly-characterized, multisite US cohort—we first identify those who initiated cannabis use through comprehensive substance use assessment, including self-report and multiple toxicological tests. We then conduct longitudinal mixed effect models to assess trajectories of neurocognitive performance in those who initiated cannabis use relative to those who did not. We hypothesize those who use cannabis will demonstrate restricted improvements and likely decline across cognitive domains, particularly in learning, memory, and working memory, based on prior longitudinal findings^10–17^. Secondarily, follow-up analyses utilize robust cannabinoid content from hair to examine whether recent use of specific cannabinoid constituents (THC v. CBD v. Controls) differentially alter neurocognitive trajectories, expecting decreased cognitive performance across domains associated with THC^37,38^, but not CBD^28,29^. Importantly, primary models covary for core confounds and primary and secondary models include within-person nesting to account for individual trajectories to account for developmental change across adolescence and neurocognition prior to cannabis use onset.

## Methods

### Participants

Participants were recruited at ages 9-10 in 2016-2018 via propensity-based school recruitment at 21 sites across the United States, resulting in a cohort of 11,868^42,43^, and followed for 10+ years. ABCD data release 6.0 contains data collected from September 1, 2016 to January 15, 2024, which includes complete annual data from Waves 0-5 and half of Wave 6. **Figure 1** displays sample sizes for the present analyses.

**Figure 1.**
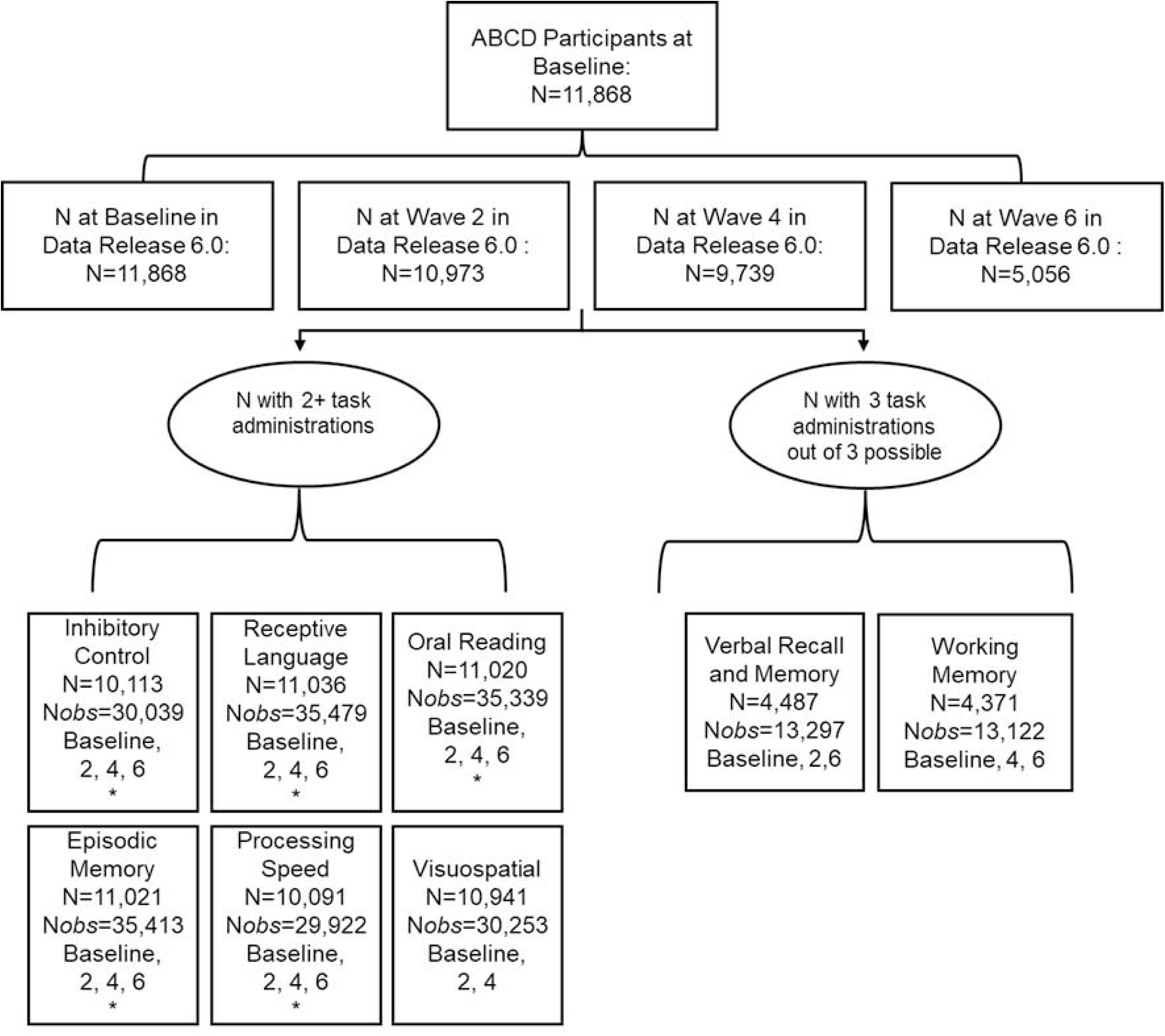
Longitudinal Sample Size by Neurocognitive Task Notes: Chart describing sample sizes and number of complete administrations required to be included in analyses. N=Sample size by task; N*obs*=number of observations included in the models. * indicates tasks that were included in secondary hair cannabinoid analyses, as these have data available at Waves 2, 4, and 6 (to match waves of available hair data); only Wave 6 data from half the cohort was released for use in the present analyses.

Eligible pasrticipants and their caregivers presented at their local site for enrollment and a Baseline session, with mid-year brief assessment and annual in-person visits each year thereafter. Annual sessions included biosample collection, self- and parent-report surveys, clinical mental health interview, and a semi-structured substance use interview^44–49^. On even years (Baseline, Wave 2, Wave 4, etc.) participants also completed a full neurocognitive battery. Starting Wave 3, all youth were told to remain substance-free 24 hours prior to the session. Participants and their caregivers completed written informed assent and consent, respectively. All study protocols were completed consistent with the Declaration of Helsinki and approved through a single IRB at the University of California, San Diego.

## Measures

### Substance Use Measures

#### Self-Reported Substance Use and Timeline Followback Interview

Prior to querying substance use, research assistants (RAs) reminded participants of confidentiality, including that caregivers will not be informed of their responses to the substance use questions unless they report clear, imminent threat or harm or other state-mandated reporting requirements.

Participants annually completed a substance use interview covering lifetime at Baseline and the past year (or since last visit) at annual follow-ups. When participants endorsed full standard substance use (1+ standard alcohol drink or any other substance use), RAs administered a Timeline Followback (TLFB) interview to collect detailed substance use patterns since their last visit (approximately past year) by each substance class^46,50^. A mid-year assessment was given at approximately the mid-point between study visits (∼6 months) covering substance use since their last visit (binary reported yes/no).

#### Medicinal CBD Use

Beginning at Wave 2, caregivers were asked, “Has your son/daughter used any CBD (or cannabidiol) products with your or your doctor’s permission, such as Epidiolex or other over the counter CBD product, in the past year?”. Youth participants and their caregivers were also asked whether they had used CBD within the past 24 hours.

#### Oral Fluid Toxicology

An oral fluid drug screen was administered to ∼5% randomly selected participants at Baseline and increased to all participants at Wave 6. Drug classes tested include amphetamines, benzodiazepines, methamphetamines, cocaine, cannabis (THC), and opioids. Oral fluid tests sensitively identify past 12-48 hours of drug use, depending on the substance^48^.

#### Urine Toxicology

At Waves 4-6, participants completed a full drug urinalysis screening for amphetamines, barbiturates, buprenorphine, benzodiazepines, cannabinoids, cocaine, MDMA, methamphetamine, methadone, opioids, oxycodone, PCP, propoxyphene, tricyclics, and cannabis (THC). Depending on drug class and frequency/amount of drug used, urine can test positive up to a month after last use.

#### Hair Toxicology

Hair was collected from all participants annually who were willing to give a sample and did not have a hair style that would be disrupted by collection (e.g., too short of hair; braids). Approximately 100mg hair 1.5cm from the root were collected from 3-4 regions around the crown of the participant’s head. Hair was securely packaged and stored until selected for testing, based on funding and prioritization of sample selection^23,51^, with testing processes and cut-offs described in the Supplement. Hair samples provide on average a 3-month window of detection for substance use^24,52,53^. Hair testing for cannabis is sensitive for detecting moderate-to-heavy use^54^ (using a minimum of twice a month on average^24^) and demonstrates excellent specificity in adolescents and young adults^24^. While some drugs may demonstrate altered binding and thus concentration of drug analytes in hair dependent on hair melanin, cannabinoids do not demonstrate this same relationship^55,56^.

Predictors of interest include: (Aim 1) <u>Any Lifetime Cannabis Use</u>, combining all cannabis data (self-report, toxicological) into a binary variable, in which all participants are coded “Controls” until any use is identified via self-reported use exceeding a puff/taste or any positive cannabis toxicology, then subsequently coded “yes" for lifetime cannabis use. Mid-year interview substance use (e.g., Wave 4.5) was applied to the following study visit (e.g., Wave 5). In models, indicators of cannabis use were entered as time-varying predictors with a binary yes/no variable representing cannabis use at the wave with first-identified use and then carried forward for each subsequent wave.

(Aim 2) <u>Hair Cannabinoids</u> coded as “Controls” (no cannabinoids detected in hair), “THC Only” (THC/THCCOOH/Δ8-THC detected only in hair), and “CBD+” (CBD detected in hair, regardless of THC/THCCOOH/Δ8-THC status) to assess the influence of recent, specific cannabinoids on neurocognition, consistent with prior work^29^; in these models, hair cannabinoid groups were time-varying at each wave. As less than monthly cannabis use is unlikely to be detected by hair testing, those who only experiment with use may be included as “Controls” when relying on hair cannabinoids alone.

## Neurocognitive Outcomes

Tasks selected for the present analyses include those given on even study years and administered at least three times between Baseline and Wave 6 (see **Figure 1** for task administration timing). Analyses were restricted to participants who completed at least two administrations of the same task across waves. For List Sort Working Memory and Rey Auditory Verbal Learning Test (RAVLT), data are only available for less than half of participants at Wave 6 and there were only three possible administrations across time. Therefore, all three administrations were required given it was not possible for a majority of participants to complete the task.

*NIH Toolbox*^57^. <u>Inhibitory control</u> was assessed via the Flanker Inhibitory Control and Attention task, where participants were to press an arrow key matching the direction a stimulus arrow was pointing (range Standard Score[SS]=41-117). For <u>receptive language</u> (Picture Vocabulary), participants listened to a word and matched it to one of four pictures (range SS=22-126). For <u>oral reading</u>, participants read and pronounced a list of words (range SS=59-180). In the List Sort <u>Working Memory</u>, participants ordered by size a list of animals and produce (range SS=36-136). Pattern Comparison <u>Processing Speed</u> asked participants to quickly identify whether two pictures were identical (range SS=30-174). Picture Sequencing Memory test assessed <u>episodic memory</u> via showing the participants a series of pictures, then asking them to sort the pictures in the same order as demonstrated (range SS=74-138). Primary outcomes included uncorrected standard scores for each subtest, consistent with prior studies to ensure meaningful results of within-participant changes^58^.

*Little Man Task*^59^. Participants were shown a “little man” holding a briefcase in various orientations to assess visuospatial processing and mental rotation. As the figure shifts (e.g., upside down; faces away from the participant), the participant must select whether the briefcase is in the man’s left or right hand. Total correct responses (range=0-32 correct items) were used as the primary <u>visuospatial</u> performance outcome.

*RAVLT*^60^. Verbal recall and memory were assessed through the RAVLT. Participants were read a list of words five times, followed by a distractor list, the original list, and then a long delay. Here, <u>immediate verbal recall</u> was assessed with number of correct Trial 1 responses (range=0-15 words) and <u>memory</u> was determined by number of correct responses after the long delay (range=0-15 words).

### Covariates

Covariates included those previously associated with cannabis and behavioral outcomes^61^; variables are described in more detail in the Supplement. Covariate use within models is described in the Statistical Analyses.

#### Time-Invariant

*Sociodemographic covariates*. Baseline sociodemographic characteristics, as reported by the caregiver, were selected as covariates given their relevance to cognition and substance use: highest level of parental education attainment^62^, biological sex of the participant as assigned at birth^63^, and self-reported race and ethnicity^45^ (a construct reflecting lived experience rather than biological categories^64^; included due to potential differences in rates of hair collection^65^). *Family History of Substance Use.* Caregivers reported whether any of the participant’s biological relatives had alcohol use problems (AUD) or, separately, substance use problems (SUD)^66^. *Prenatal Substance Exposure*. Caregivers reported whether the participant was exposed to any drug classes in utero, both before and after knowledge of the pregnancy^45^.

Each drug class was coded as its own exposure (1/0, regardless of whether using before/after knowledge of pregnancy; see Supplement for details). *Baseline Psychopathology*. Caregivers completed the Childhood Behavior Checklist (CBCL), with age- and sex-normed t-scores subscales for Externalizing and Internalizing symptoms included as covariates, given youth psychopathology is often identified as a predictor of substance initiation^67^.

#### Time-Varying

Youth age at each Wave was included. *Other Substance Use*. Polysubstance use is common and may influence cognition^68,69^. Using the same identification method of lifetime substance exposure as described for cannabis use, binary (yes/no) time-varying lifetime alcohol use, nicotine use, and any other substance use variables were computed across all observations.

## Analytic Plan

ABCD Data Release 6.0 was used (https://doi.org/10.82525/jy7n-g441). Analyses were conducted in R. Random intercepts for participant, family, and site were used. Models allowing random age slopes were evaluated but did not improve fit and produced convergence warnings; therefore, random intercept models were retained.

### Aim 1: Longitudinal Influence of Cannabis Onset

Linear mixed effect models were run using *lme4* package in R^70^. Neurocognitive task performance over time was predicted by lifetime cannabis use group status (CU group) interacting with age. Non-linear, quadratic models were examined for all cognitive measures; inclusion of age² terms did not improve model fit (all ΔAIC > 2; all p > .10). The one exception was for Picture Vocabulary, wherein the quadratic interaction was significant and improved fit. For parsimony, only linear models are presented here for all cognitive tasks with the Picture Vocabulary quadratic model presented in the Supplement.

Covariates of interest, often time-varying, were included in the full linear mixed effect models: lifetime alcohol use, lifetime nicotine use, lifetime other substance use, sex, and age. Additionally, for variables shown to be linked to cannabis group status and cognition which are more static in nature or potential confounds of no interest (i.e., prenatal exposure, family history of substance use disorder, baseline psychopathology, sociodemographic factors), an inverse propensity score was created to account for covariates while also reducing variable dimensionality (see Supplement for specific variables and **Figure S2** for evidence of balance). To estimate the inverse propensity score variable, a logistic regression was run, predicting lifetime cannabis exposure by all listed factors, clustering on site and family ID. The inverse of logistic regression values (their individual probability of exposure) were saved for each participant and included as a covariate in analyses.

Study site, family ID, and subject ID (to account for within-person change) were included as nested effects. FDR corrections were applied within models^71^. Standardized β were calculated using the *MuMin* package^72^, acknowledging that ABCD is powered to detect very small effect sizes^73^. Estimated marginal means by age were extracted via the *emmeans* package^74^. 95% Wald confidence intervals are included. Interpretation of significance was set at FDR-*p*<.05 for primary models.

### Aim 2: Longitudinal Influence of Hair Cannabinoids

To specifically assess potential recent THC or CBD influence relative to no use, a subsample of the cohort was included in secondary longitudinal analyses. All participants with hair tested at Waves 2, 4, and 6 were included (n=648). Time-varying groups by wave were then defined by hair cannabinoid content: no cannabinoids present (Controls n=546); THC (identified by THC or THCCOOH; THC Only Group n=81); and CBD (identified by confirmed CBD, regardless of whether THC/THCCOOH were present; CBD+ group n=21), consistent with prior research^29^. Notably, n=13 in the CBD+ group also had THC present in their hair sample and n<10 participants with hair cannabinoids were both in the CBD and THC groups on altering years, though descriptively are included within the CBD group (**Table S2**). Mean performance by hair cannabinoid group and age are presented in **Table S3**.

Linear mixed effects models were run to predict cognitive performance by hair group*age. Analyses included tasks that were administered at the same waves as selected hair samples. Given limited prior research using novel repeat hair testing results and cognition and due to the smaller cell sizes, use of covariates were constrained to more sensitively assess potential cannabinoid relationships: sex assigned at birth and any other substance use (i.e., alcohol, nicotine, or any other substance combined) positive in hair. Models were nested for family ID and subject ID; study site was included as a fixed effect when convergence issues were identified. Interpretation of significance was set at *p*<.01 for hair cannabinoid models.

## Results

### Participant Sociodemographic and Substance Use Characteristics

Participants included the full ABCD cohort (see **Table 1** for descriptives). Those who used cannabis by the Wave 6 follow-up differed from those who did not in sex (χ^2^=7.3, *p*=.007), race (χ^2^=62.9, *p*<.0001), and parental educational attainment (χ^2^=147.3, *p*<.0001). Mean group scores on externalizing and internalizing symptoms by wave are also presented in **Table S3**. Onset of cannabis use was most commonly identified by self-report (67%), followed by hair testing (22%), urine toxicology (21%), oral fluid testing (5%), and self-/parent-reported medicinal CBD use (3%; percentages not mutually exclusive) (i.e., some participants were identified solely by self-report while others were identified through toxicological tests).

**Table 1.**
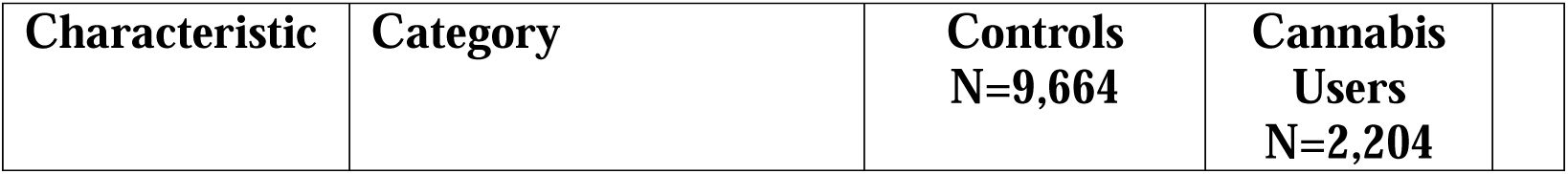
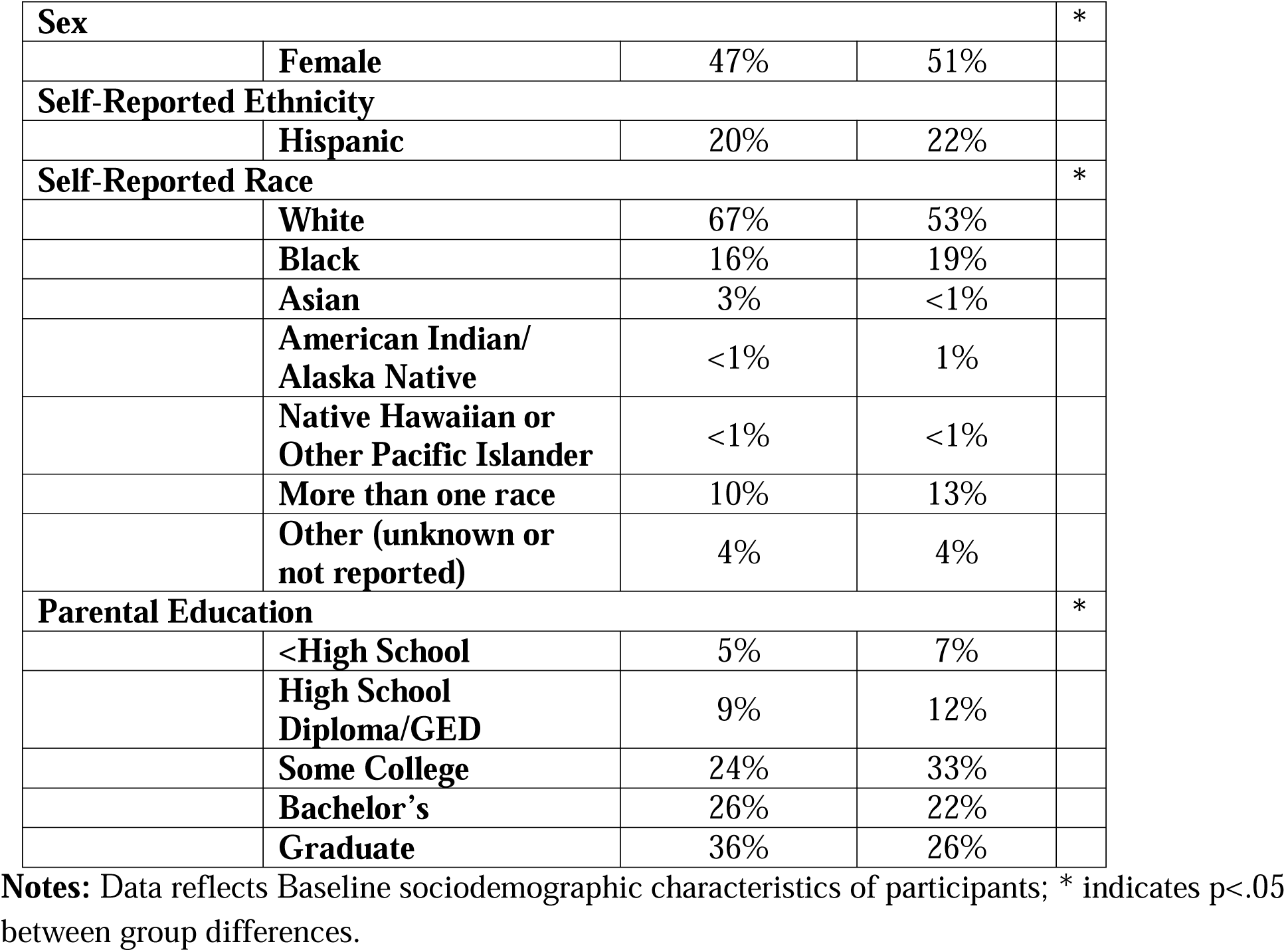
Sociodemographics of groups

| Characteristic | Category | Controls<br>N=9,664 | Cannabis<br>Users<br>N=2,204 |
| --- | --- | --- | --- |

|  |  |  |  |  |
| --- | --- | --- | --- | --- |
| **Sex** |  |  |  | * |
|  | **Female** | 47% | 51% |  |
| **Self-Reported Ethnicity** |  |  |  |  |
|  | **Hispanic** | 20% | 22% |  |
| **Self-Reported Race** |  |  |  | * |
|  | **White** | 67% | 53% |  |
|  | **Black** | 16% | 19% |  |
|  | **Asian** | 3% | <1% |  |
|  | **American Indian/**<br>**Alaska Native** | <1% | 1% |  |
|  | **Native Hawaiian or**<br>**Other Pacific Islander** | <1% | <1% |  |
|  | **More than one race** | 10% | 13% |  |
|  | **Other (unknown or**<br>**not reported)** | 4% | 4% |  |
| **Parental Education** |  |  |  | * |
|  | **<High School** | 5% | 7% |  |
|  | **High School**<br>**Diploma/GED** | 9% | 12% |  |
|  | **Some College** | 24% | 33% |  |
|  | **Bachelor's** | 26% | 22% |  |
|  | **Graduate** | 36% | 26% |  |

**Notes:** Data reflects Baseline sociodemographic characteristics of participants; \* indicates $p < .05$ between group differences.

For secondary models, hair groups (THC, CBD, or Controls) differed by sex (χ^2^=12.4, *p*=.002), parental education (χ^2^=64.8, *p*<.0001), race (χ^2^=46.9, *p*<.0001), and ethnicity (χ^2^=13.1, *p*=.001), with descriptives included in **Table S4**.

### Aim 1: Primary Neurocognitive Models

As aims were to investigate longitudinal trajectories of cognition, only the interaction between CU status and age are reported, though main effects were found across tasks (FDR-*p*s<.05). Full models are included in **Table S5** and mean performances by group and age are in **Table S6**; mean group differences are presented at *p*<.05.

*Working Memory:* Working memory performance exhibited a significant CU*age interaction with medium effect size (β=-0.52, b=-1.32, 95%CI:-1.80,-0.84, *p*<.0001, FDR-*p*<.0001), such that CU participants did not demonstrate the same improvement in working memory with age as Controls. Review of estimated marginal means indicates groups differed from ages 9-14 with CU group demonstrating better performance (*p*s<.0001), whereas at 15-16 groups did not differ (**Figure 2a**). At age 17, CU exhibited toward worse performance (*p*=.0001).

**Figure 2.**
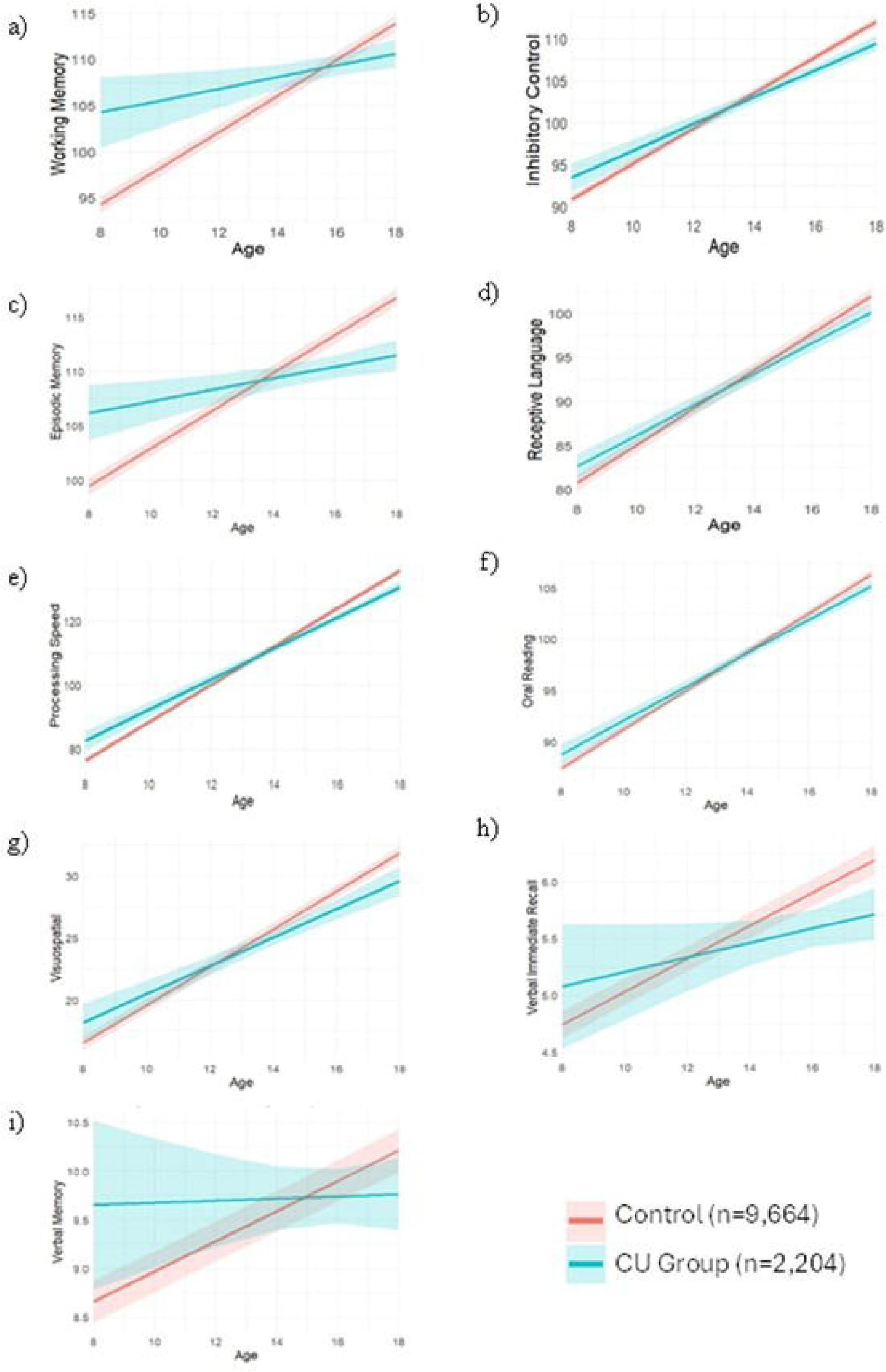
Cannabis Group Status * Age on Neurocognitive Performance Notes: Cannabis use group status is determined by time-varying lifetime cannabis use, where once cannabis use is identified that participant is always included in the cannabis use (CU) group. Cannabis use group status interactions with age by neurocognitive domain; 2a-f y-axes reflect uncorrected NIH Toolbox performance by domain [working memory n=4,371; picture memory n=11,021; vocabulary n=11,036; processing speed n=10,091]; 2g is the Little Man Task number correct scores [n=10,941]; 2h and 2i are Rey Auditory Verbal Learning Test (RAVLT) immediate recall (Trial 1) and memory (delayed recall) raw scores, respectively [n=4,487]. Total sum N of those who used cannabis are in blue (n=2,204); those who did not use cannabis are in red (n=9,664). Groups were time-varying with onset identified over time.

*Inhibitory Control*: Inhibitory control exhibited a significant CU*age interaction with small effect size (β=-0.21, b=-0.52, 95%CI:-0.73,-0.31, *p*<.0001, FDR-*p*<.0001; **Figure 2b**). The CU group performed better at ages 9-10 (*p*s<.01) and 11 (*p*=.03), with the relationship flipping starting at age 14 (*p*=.03) through 15-17 (*p*s<.0001).

*Episodic Memory:* CU group interacted with age with medium effect size (β=-0.32, b=-1.20, 95%CI:-1.54,-0.86, *p*<.0001, FDR-*p*<.0001; **Figure 2c**) in predicting episodic memory. At ages 9-13, CU demonstrated better episodic memory performance (*p*s<.01), no difference at ages 13-14, and worse performance than Controls at ages 15-17 (*p*s<.001).

*Receptive Language*: There was an interaction of CU*age in receptive language with small effect size (β=-0.14, b=-0.36, 95%CI:-0.51,-0.21, *p*<.0001, FDR-*p*<.0001), with the CU group demonstrating less change over time (**Figure 2d**). At ages 9-10, CU demonstrated better performance (*p*s<.01), with worse performance at ages 14 (*p*=.03) and 15+ (*p*s<.0001). Follow-up analyses indicated a quadratic relationship (CU*age^2^) predicting performance, which is detailed in the Supplement.

*Processing Speed:* The interaction of CU*age (β=-0.24, b=-1.13, 95%CI:-1.53,-0.74, *p*<.0001, FDR-*p*<.0001) significantly predicted processing speed with small effect size, with Controls demonstrating greater improvement with age (**Figure 2e**). At ages 9-12, CU demonstrated better processing speed (*p*s<.05), with no difference ages 13-14, and CU demonstrating worse performance than Controls at ages 15-17 (*p*s<.0001).

*Oral Reading:* For oral reading, there was an interaction of CU*age with small effect size (β=-0. 11, b=-0.25, 95%CI:-0.37,-0.12, *p*=.0001, FDR-*p=*.0003; **Figure 2f**). CU group performed better at ages 9-11 (*p*s<.05) and worse than Controls at ages 15-17 (*p*s<.01).

*Visuospatial:* For predicting number correct on visuospatial performance, site was included as a fixed effect due to convergence issues. Results indicate a significant interaction between CU*age with small effect size (β=-0.16, b=-0.39, 95%CI:-0.63,-0.16, *p*=.001, FDR-*p*=.003), such that those in the CU group demonstrated lower number correct with age (**Figure 2g**). While CU exhibited more correct responses than Controls at age 9 (*p*=.04) and no difference at ages 10-13, Controls had more correct responses at ages 15-17 (*p<*.0001).

*Verbal Recall and Memory:* CU group*age significantly interacted to predict immediate recall with small effect size (β=-0.21, b=-0.08, 95%CI:-0.15,-0.01, *p*=.02, FDR-*p*=.03; **Figure 2h**). Groups did not differ from ages 9-14, though in ages 15-17 the CU group demonstrated worse performance than Controls (*p*s<.05). For long delay memory, there was also a significant interaction between CU*age (β=-0.21, b=-0.14, 95%CI:-0.25,-0.04, *p*=.008, FDR-*p*=.01; **Figure 2i**), such that those in the CU group demonstrated less improvement over time. Groups differed from ages 9-11 with CU group demonstrating better performance (*p*s<.05), whereas at 12-16 there was no difference between groups, and at age 17 CU demonstrated worse performance (*p*=.02).

### Aim 2: Secondary Hair Analyses

After controlling for sex, other positive hair toxicology results, and site, and nesting for family ID and subject ID, THC Only group status*age significantly predicted <u>episodic memory</u> performance relative to Controls with medium effect size (β=-0.60, b=-2.37, 95%CI:-4.08,-0.66, *p*=.007; n=645, n_observations_=1,885; see full models in **Table S7**). Higher scores reflect better episodic memory performance; thus, the negative THC*age interaction indicates a reduced rate of age-related improvement among THC-positive youth (**Figure 3**). Estimated marginal means revealed that at ages 15-17, THC demonstrated worse episodic memory than Controls (*p*s=.04-

**Figure 3.**
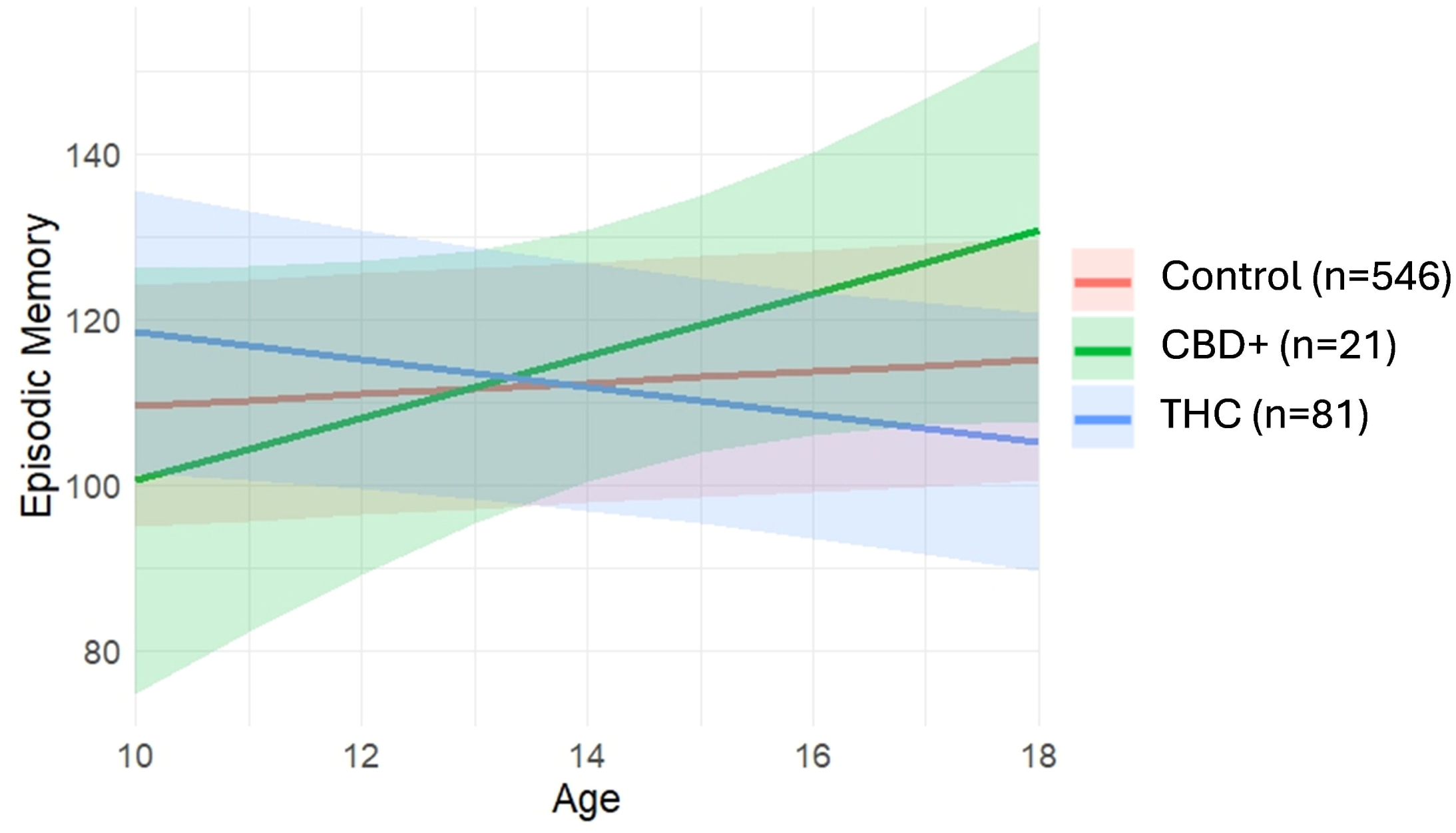
Hair Cannabinoid Group*Age on Episodic Memory Notes: Hair cannabinoid use group identified by whether there was THC (without CBD) in hair, CBD in hair (regardless of THC), or no hair cannabinoids. Hair testing and episodic memory (NIH Toolbox Picture Memory Uncorrected Standard Score) testing occurred every other year (approximately ages 11-12, 13-14, and 15-16). Total sum N of those without hair cannabinoids (Controls) are in red (n=546); those with CBD in hair are in green (n=21); those with THC are in blue (n=81). Groups were time-varying with onset identified over time.

.001) and worse performance than CBD+ (*p*s=.02-008). CBD+ group and Controls were not significantly different. Given the relatively small CBD-positive subsample (n=21), these analyses should be interpreted cautiously. For all other neurocognitive tasks (receptive language, processing speed, inhibitory control, and oral reading), no significant relationships were found between hair cannabinoid groups and performance.

## Discussion

Findings support and extend prior work suggesting aberrant neurocognitive developmental trajectories in youth ages 9-17 who use cannabis, significantly extending the existing literature by including a comprehensive assessment of self-report and objective toxicological measures to accurately categorize cannabis use onset. Utilizing a prospective longitudinal design that considered pre-substance use confounding factors (i.e., sociodemographics, prenatal substance use history, family history of AUD/SUD, internalizing and externalizing symptoms) and controlled for comorbid substance use onset, we found that those who use cannabis actually show a slight cognitive advantage during late childhood, but demonstrate restricted improvement trajectories in immediate recall and delayed memory, processing speed, inhibitory control, visuospatial processing, language, and working memory over adolescence. Follow-up analyses considered a potential mechanism of altered trajectories through unique contributions of THC and CBD in analyzed in objective hair toxicology. Those with THC use exhibited declining trajectories in episodic memory relative to those without hair cannabinoids, whereas those with CBD did not differ from controls.

In the present study the cannabis use group demonstrated either restricted improvement or plateaued performance relative to non-using youth across cognitive domains. Longitudinal studies previously identified changes in those who initiated cannabis use in learning and memory^10–16,18^, working memory^11,12,16,17^, inhibition and attention^12,15,16,18^ visuospatial functioning^11,12,14,16^, and processing speed^10,12,14,15^. Here, those who use cannabis exhibited potential neurocognitive advantage in earlier years with a diminished positive slope over time. While on most tasks both groups demonstrated improvement over time as expected^58^, those who used cannabis did not demonstrate the same rate of improvement. It may be that cannabis itself induces neurotoxicity which in turn impacts cognitive trajectories, or there may be a shared vulnerability for both cannabis use and reduced gains in cognition over time^75^.

Other prior work within this cohort describes higher cognition being associated with substance use onset and cannabis specifically^76,77^. It may be that early maturation increases risk of using, and that these youth then demonstrate restricted age-typical cognitive improvement over time. Demonstrated cognitive trajectories may also reflect regression to the mean. Other analyses suggest family and environmental factors may most robustly predict substance initiation^76^, and therefore broader consideration of such factors (e.g., personality traits such as openness to experience^78,79^) should be considered in future analyses. Following this cohort into young adulthood can determine how variations in use trajectories, including escalation of use and cessation, influence cognition over time.

Secondary analyses indicate that youth who have objective evidence of recent THC, but no CBD, exposure in hair samples are likely to demonstrate worse episodic memory performance over time than those with no cannabinoids in their hair. Importantly, as hair cannabinoid testing identifies more frequent cannabis use than experimentation, these secondary models may also reflect more heavy cannabis use than the primary models. Findings align with prior hair cannabinoid results in adults^29^, and are generally consistent with cross-sectional analyses of CBD hair cannabinoids in this cohort that did not reveal CBD-cognition relationships^25^. These data indicate a potential mechanism (THC) driving the displayed neurocognitive trajectories observed in the larger CU analyses, consistent with theorized processes^27,80^. THC, then, is likely particularly important for otherwise healthy adolescents to avoid or delay use. On balance, other data within this cohort indicated that even those who use CBD for medicinal reasons may be unintentionally exposed to THC^81^, and product mislabeling is a known concern even in regulated markets^82–85^. Hair cannabinoid models also included far fewer participants and the sample size for the CBD group was low (n=21), limiting power and interpretation. We caution that null findings in CBD models should not be interpreted as evidence of absence of association.

Therefore, while CBD may be considered a harm reduction option for those regularly using cannabis in the context of mitigating THC-related consequences or cessation, the exact impact of CBD and the potential for contamination of CBD (including with THC) remain significant concerns for adolescent health.

Reasons for variation in findings between primary and secondary analyses warrant consideration. First, a smaller sample size with fewer waves of data incorporated may explain some discrepancy due to reduced power in hair cannabinoid models. In addition, modeling techniques were substantially different, including different cannabis use identification methods (comprehensive measures of *any* use in lifetime v. recent hair cannabinoid content alone) and different sets of covariates. Primary models may then reflect a more shared vulnerability for neurocognition that reflects both cannabis use and other confounds, while hair cannabinoid models may indicate a THC-specific vulnerability in episodic memory.

Small-to-medium small effect sizes were observed, comparable to prior research^86^ and consistent with most data analyses of this size^73^. Importantly, given the sensitive sociodevelopmental period of adolescence, even modest effects may translate to clinically significant alterations^87,88^. For instance, relative decrements in learning and memory can impact standardized testing, grade advancement, and higher education opportunities in a highly competitive contemporary environment. Similarly, visuospatial, processing speed, and working memory skills are paramount to driving, an area where teens may already be at a disadvantage^89,90^. The impact of these altered neurocognitive trajectories may be meaningful in individual lives, particularly when individuals initiate use before age 16^91^, though further investigation to the real-world impact of such cannabis use is needed.

Findings are considered in light of strengths and limitations. Results here expand on the prior literature through several notable methodological improvements: (1) use of subjective (self-report) and objective (toxicological) cannabis use variables; (2) wide consideration of potential confounds, including prenatal history, family factors, sociodemographics, other substance use, and within-person change; and (3) a large cohort that was almost exclusively substance naïve at Baseline (ages 9-10)^92^, with neurocognitive measures that predate cannabis exposure. On balance, secondary analyses of THC/CBD consisted of a smaller subsample, including a relatively small number of those with CBD in their hair. Motivations for cannabis and potentially CBD use are not known, though may include efforts to treat underlying symptoms (e.g., sleep, pain, anxiety) which may confound differences from healthy controls. Residual time-varying confounding cannot be excluded. Although false discovery rate correction was applied in primary models, testing across multiple outcomes and smaller subsamples increases risk of both Type I and Type II error. Oral fluid and urinalysis tests were not confirmed, though both demonstrate adequate specificity^24,36^ and hair samples only positive for THC or CBD may be related to external contamination^93^; inclusion of toxicological measures here are still more likely to contribute to more accurate grouping rather than mis-categorizing someone as using cannabis when they do not. Hair samples were tested in two different laboratories with their own methods and LOD/LOQ/cutoffs, which increases noise in hair cannabinoid interpretation. While efforts were made to recruit a diverse cohort^42^, there are still selection biases that may reduce generalizability. Participants included here are largely in the middle of adolescence and will undergo more neurocognitive development; early cannabis use (e.g., before age 16) is known to be particularly risky for adolescents^91^ and future analyses of ABCD should include consideration of age of onset of regular use. Additional follow-up to consider trajectories of those who initiate cannabis use or continue to use in later adolescence, including modeling of frequency of use, are needed.

## Conclusions

Adolescence marks a period of substantial development, but cannabis use may constrain one’s full cognitive developmental performance. Data here indicate novel restricted improvement and flattened neurocognitive trajectories in youth (ages 9-17) who initiate cannabis use, after accounting for within-person change and numerous known confounds and improving accuracy in cannabis groupings through incorporating toxicological measures. Unique analyses harnessing objective toxicology demonstrate that THC use may specifically relate to episodic memory performance over time during this developmental period. Findings support interventions aimed at delaying cannabis initiation during early adolescence and integrating neuroscience-informed psychoeducation about cognitive development during sensitive periods^94–96^. Continued monitoring of this cohort will clarify cannabinoid-cognition relationships into young adulthood, including the impact of timing of cannabis use initiation.

## Acknowledgements

Authors would like to thank the families who participate in the ABCD Study and the staff who collect the data. Data used in the preparation of this article were obtained from the <u>Adolescent Brain Cognitive Development™ (ABCD) Study</u>, held in the <u>NIH Brain</u> <u>Development Cohorts Data Sharing Platform</u>. This is a multisite, longitudinal study designed to recruit more than 10,000 children aged 9–10 and follow them over 10 years into early adulthood.

ABCD Consortium investigators designed and implemented the study and/or provided data but did not necessarily participate in the analysis or writing of this report. This manuscript reflects the views of the authors and may not reflect the opinions or views of the NIH or ABCD Consortium investigators. The ABCD data repository grows and changes over time. The ABCD data used in this report came from https://doi.org/10.82525/jy7n-g441. Authors can be contacted for analytical code.

## Author Contributions

NEW conceptualized, conducted statistical analyses, interpreted fndings, drafted the original manuscript, and completed critical revisions. RMS and ALW assisted in conceptualization, data analysis, interpretation, and critical revisions. RV assisted in design, stastisical methods, and critical revisions. VS, KML, MAH, PDB, HB, LM, and JJ all contributed to conceptualization, interpretation, and critical revisions. SFT contributed to conceptualization and design, interpretation, and critical revisions. All authors contributed to the writing and revision of the manuscript and have reviewed and approved the final version of the manuscript.

## Funding

N.E. Wade was supported by National Institute on Drug Abuse (DA050779, PI: Wade). This work was also supported by DA064409 (PI: Sullivan), DA062011 (PI: Wallace), T32 AA013525 (PI: Riley/Spadoni to Szpak), R01DA062432 (MPI Lisdahl, Hillard), 2U01DA041025 (MPI Lisdahl, Larson), K01DA057389 (PI: Gonçalves), and NARSAD/Brain Behavior Research Foundation (Gonçalves).

The ABCD Study® is supported by the **National Institutes of Health** and additional federal partners under award numbers: U01DA041048, U01DA050989, U01DA051016, U01DA041022, U01DA051018, U01DA051037, U01DA050987, U01DA041174, U01DA041106, U01DA041117, U01DA041028, U01DA041134, U01DA050988, U01DA051039, U01DA041156, U01DA041025, U01DA041120, U01DA051038, U01DA041148, U01DA041093, U01DA041089, U24DA041123, U24DA041147.

A full list of supporters is available at Federal Partners – ABCD Study.

Additional support for this work was made possible from NIEHS R01-ES032295 and R01-ES031074.

## Competing Interests

Authors have no conflicts of interest to disclose.

## Notes

https://www.nbdc-datahub.org/

### Competing Interest Statement

The authors have declared no competing interest.

### Summary of Updates

-Introduction rationale clarified -Methods include more details (including in the supplement) -Supplemental analyses conducted, full models included in supplement -Discussion expanded

