## Supplement for "Longitudinal Neurocognitive Trajectories in a Large Cohort of Youth Who Use Cannabis: Combining Self-Report and Toxicology"

**Supplemental Materials**

Hair Testing Description	p. 2

Table S1. LOQ/LOD by Lab

Figure S1. Love Plot Demonstrating Covariate Balance Before/After Adjustment

Covariate Descriptions and Inverse Propensity Score Calculation	p. 3

Non-Linear Cannabis-Cognition Relationships	p. 6

Table S2. Quadratic Receptive Language Model

Figure S2. Quadratic, Non-Linear Receptive Language Trajectories

Table S3. Mean CBCL Externalizing and Internalizing Symptoms by Group and Wave	p. 8

Table S4. Sociodemographics within Hair Analyses	p. 9

Table S5. Primary Neurocognitive Models Full Output	p. 10

Table S6. Mean Performance on Cognitive Tasks by Group and Age	p. 20

Table S7. Secondary Neurocognition-Hair Group Models Full Output	p. 23

Table S8. Mean Performance on Neurocognitive Tasks by Hair Cannabinoid Group	p. 31

and Age

**Hair Testing Description**

Selected hair samples were shipped to Psychemedics (Culver City, CA) or USDTL (Des Plaines, IL) for testing (see Wade et al., preprint, https://doi.org/10.1101/2025.09.19.25336190). Hair underwent screening for possible drug exposure. Presumptive positive samples were washed to remove external contamination^1^, though secondhand exposure cannot be ruled out in samples only positive for THC or CBD. Presumptive positives samples were then confirmed by LC/MS-MS or GC/MS-MS testing, depending on the drug class. Hair samples provide specific information on parent cannabinoids (i.e., ∆9-THC, CBD, ∆8-THC, ∆10-THC) and the primary THC metabolites (THCCOOH), which confirm THC ingestion^2^. Limits of quantification (LOQ), limits of detection (LOD), and cut-offs for cannabinoids in hair in testing laboratories are in **Table S1.**

**Table S1.** LOQ/LOD by Lab

| **Psychemedics** | |  |  |  | **USDTL** | |  |  |  |
| --- | --- | --- | --- | --- | --- | --- | --- | --- | --- |
| **Drug Class** | **Analyte** | **LOD/LOQ** | **Cutoff** |  | **Drug Class** | **Analyte** | **LOD** | **LOQ** | **Cutoff** |
| **Nicotine** | | |  |  | **Nicotine** | |  |  |  |
|  | Nicotine | -- | -- |  |  | Nicotine | 20 | 40 | 100 |
|  | Cotinine | 50 | 50 |  |  | Cotinine | 20 | 40 | 100 |
| **Parent Cannabinoids** | | |  |  | **Parent Cannabinoids** | | |  |  |
|  | THC | 5 | 5 |  |  | THC | 40 | 40 | 40 |
|  | CBN | 5 | 5 |  |  | CBN | -- | -- | -- |
|  | CBD | 5 | 5 |  |  | CBD | 40 | 40 | 40 |
|  | THCV | 5 | 5 |  |  | THCV | -- | -- | -- |
|  | Delta-8-THC | -- | -- |  |  | Delta-8-THC | 40 | 40 | 40 |
|  | Delta-10-THC | -- | -- |  |  | Delta-10-THC | 40 | 40 | 40 |
| **THCCOOH** | | |  |  | **THCCOOH** | |  |  |  |
|  | THCCOOH | 0.02 | 0.02 |  |  | THCCOOH | 0.01 | 0.02 | 0.05 |
| **Alcohol** | | |  |  | **Ethyl Glucuronide** | |  |  |  |
|  | Ethyl Glucuronide | 1 | 1 |  |  | Ethyl Glucuronide | 4 | 8 | 20 |

Notes: Cutoffs, LOQ, LOD are in picograms/milligram

**Covariate Descriptions and Inverse Propensity Score Calculation**

**Other substance use.** For calculating lifetime other substance use to be used as covariates, the same methods were used as in the lifetime cannabis variable. Specifically: for alcohol, any self-reported use was combined with breathalyzer and hair testing results; for nicotine, self-reported use was combined with urine and hair results; due to low identification rates, all other drug classes were combined into one “other” substance use variable. Binary lifetime use was time-varying as with lifetime cannabis use.

**Inverse Propensity Score**. Covariates within the primary neurocognitive models included both specific variables (lifetime alcohol use, lifetime nicotine use, lifetime other substance use, sex, and age, as well as nested variables for participant ID, study site, and family ID) as well as an inverse propensity score (IPS) covariate. The creation of the IPS was to both account for other common confounds while using a dimension reduction strategy. Variables included in the IPS are linked to cannabis group status and cognition and are more static in nature or potential confounds of no interest (i.e., prenatal exposure, family history of substance use disorder, baseline psychopathology, sociodemographic factors). Specifically, variables included:

- Any biological family history of alcohol problems (1/0)
- Any biological family history of drug use problems (1/0)
- Prenatal history of alcohol exposure, regardless of mother’s knowledge of pregnancy (1/0)
- Prenatal history of cocaine exposure, regardless of mother’s knowledge of pregnancy (1/0)
- Prenatal history of cannabis exposure, regardless of mother’s knowledge of pregnancy (1/0)
- Prenatal history of nicotine exposure, regardless of mother’s knowledge of pregnancy (1/0)
- Prenatal history of opiate exposure, regardless of mother’s knowledge of pregnancy (1/0)
- Prenatal history of prescription pain medication exposure, regardless of mother’s knowledge of pregnancy (1/0)
- Prenatal history of other drug exposure, regardless of mother’s knowledge of pregnancy (1/0)
- Parent-reported CBCL externalizing symptoms t-score at Wave 0
- Parent-reported CBCL internalizing symptoms t-score at Wave 0
- Ethnicity
- Race
- Highest level of parental education
- Site (nested)
- Family ID (nested)

In order to run the logistic regression to calculate inverse propensity scores, complete cases were required. Therefore, multiple imputations via the MICE package^3^ were used on missing data to enable propensity score calculation: prenatal substance use exposure (<0.1% missingness for each drug class), family history of AUD (1.2%) or SUD (1.2%), baseline internalizing (<0.1%) and externalizing symptoms (<0.1%), race (<0.1%), ethnicity (0.4%), and parental education (0.1%).

To estimate the inverse propensity score variable, logistic regression was run, predicting lifetime cannabis exposure by all listed factors, clustering on site and family ID. The inverse of logistic regression values (their individual probability of exposure) were saved for each participant and included as a covariate in analyses. Saved scores were normally distributed, with skew introduced if trimmed (e.g., at 95^th^ percentile); therefore, weights were left untrimmed. Balance of the resulting propensity score was excellent, as demonstrated in the following Love plot (**Figure S1**):

**Figure S1. Love Plot Demonstrating Covariate Balance Before/After Adjustment**

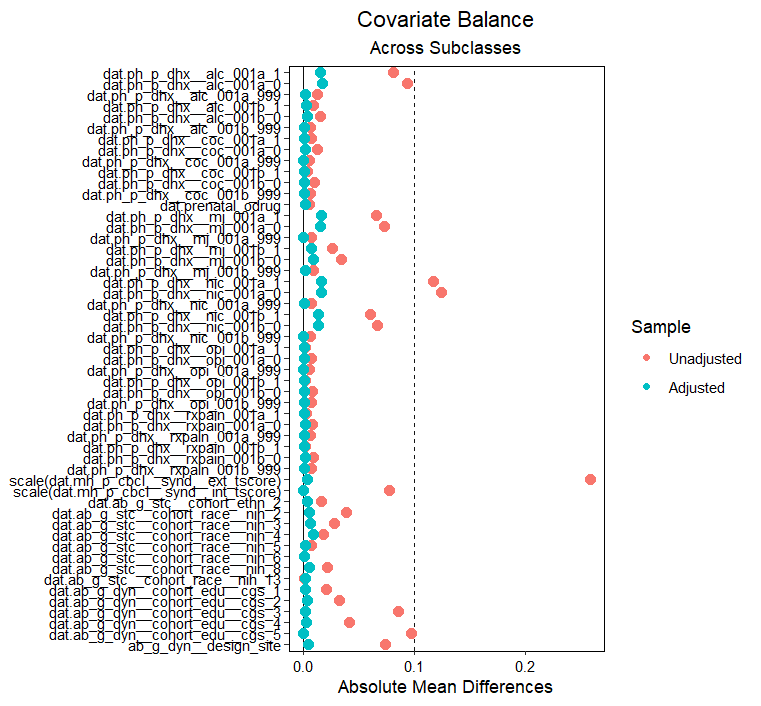

**Non-Linear Cannabis-Cognition Relationships**

Neurocognitive models were assessed for linear versus non-linear fit. Quadratic age terms were added to all primary models. Inclusion of age² terms did not improve model fit (all ΔAIC > 2; all p > .10), and linear models were retained for parsimony. The one exception is for Picture Vocabulary, where the quadratic interaction was significant and improved fit (AIC quadratic = 228,167; AIC linear = 228,343).

In the quadratic model, both CU*age (β=0.92, *p*=.019, FDR-*p*=.023) and CU*age^2^ (β=-0.51, *p*=.015, FDR-*p*=.023) exhibited significant interactions in predicting Receptive Language performance (see **Table S2**). While in general participants demonstrated improvement in Picture Vocabulary performance at earlier ages, plateauing over time, cannabis using youth demonstrated reduced improvement at an earlier age than Controls (see **Figure S2**).

**Table S2. Quadratic Receptive Language Model**

| **Receptive Language** | **Picture Vocabulary Task (NIH Toolbox): Uncorrected Standard Score** | | |
| --- | --- | --- | --- |
| *Predictors* | *Estimates* | *CI* | *FDR-p* |
| (Intercept) | 49.74 | 47.77 – 51.70 | **<0.001** |
| CU Group | -17.01 | -31.41 – -2.60 | **.029** |
| Age | 3.86 | 3.59 – 4.13 | **<0.001** |
| Age^2^ | -0.07 | -0.08 – -0.06 | **<0.001** |
| canvalue_iw | 0.91 | 0.77 – 1.04 | **<0.001** |
| any_nic_lft_yn | -0.86 | -1.17 – -0.55 | **<0.001** |
| any_alc_lft_yn | 0.19 | -0.11 – 0.50 | .219 |
| any_osu_lft_yn | -0.35 | -0.64 – -0.06 | **.029** |
| Cohort description: Participant's sex: Female | -0.47 | -0.72 – -0.21 | **<0.001** |
| CU Group * Age | 2.39 | 0.40 – 4.39 | **.029** |
| CU Group * Age^2^ | -0.09 | -0.15 – -0.02 | **.029** |
| **Random Effects** | | | |
| σ^2^ | 19.30 | | |
| τ_00_ _participant_id_ | 10.27 | | |
| τ_00_ _ab_g_stc__design_id__fam_ | 35.70 | | |
| τ_00_ _ab_g_dyn__design_site_ | 4.04 | | |
| ICC | 0.72 | | |
| N _participant_id_ | 11036 | | |
| N _ab_g_dyn__design_site_ | 22 | | |
| N _ab_g_stc__design_id__fam_ | 9120 | | |
| Observations | 35479 | | |
| Marginal R^2^ / Conditional R^2^ | 0.241 / 0.789 | | |

**Figure S2. Quadratic, Non-Linear Receptive Language** **Trajectories**

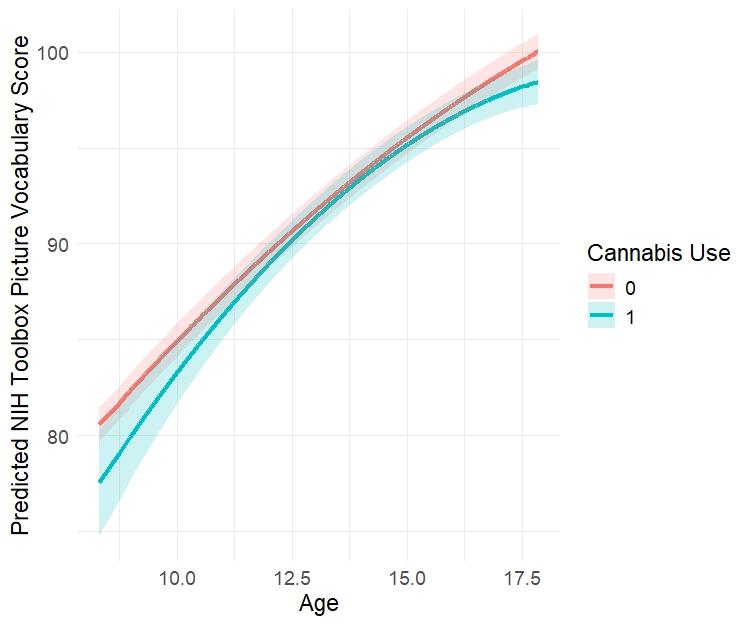

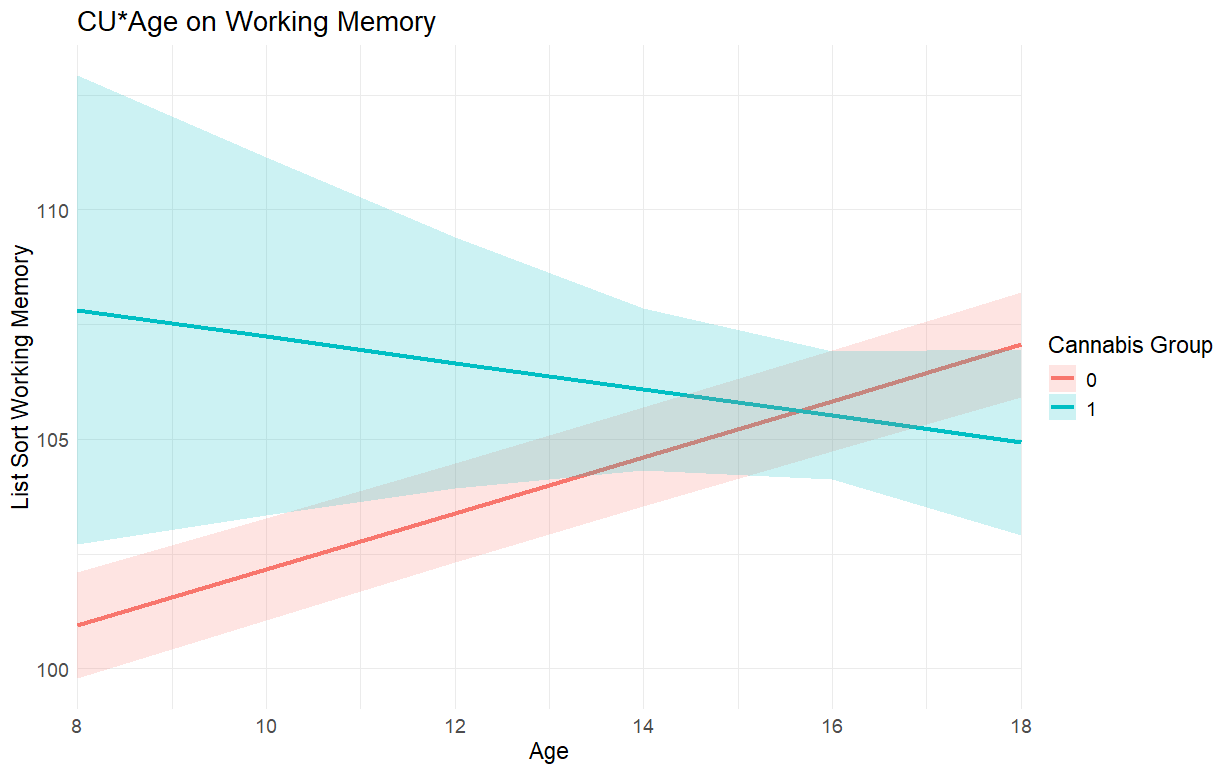

Control (n=9,058)

CU Group (n=1,978)

Notes: Cannabis use group status is determined by time-varying lifetime cannabis use, where once cannabis use is identified that participant is always included in the cannabis use (CU) group. Cannabis use group status interactions with age^2^ by Picture Vocabulary Performance (Receptive Language).

**Table S3. Mean CBCL Externalizing and Internalizing Symptoms by Group and Wave**

| **Externalizing Symptoms** | **Controls** | **CU Group** |
| --- | --- | --- |
| **Wave 0** | 45.7 (10.33) | 54.5 (11.47) |
| **Wave 2** | 44.3 (9.76) | 49.6 (12.02) |
| **Wave 4** | 43.1 (9.11) | 48.7 (11.41) |
| **Wave 6** | 41.4 (8.24) | 46.7 (10.60) |
| **Internalizing Symptoms** | **Controls** | **CU Group** |
| **Wave 0** | 48.45 (10.64) | 51.95 (11.86) |
| **Wave 2** | 47.6 (10.52) | 50.5 (11.02) |
| **Wave 4** | 47.3 (10.76) | 50.1 (11.98) |
| **Wave 6** | 46.6 (10.65) | 48.6 (11.47) |

**Table S4. Sociodemographics and Other Characteristics within Hair Analyses**

| **Characteristic** | **Category** | **Controls**  **N=546** | **THC Only Group**  **N=81** | **CBD+ Group**  **N=21** |  |
| --- | --- | --- | --- | --- | --- |
| **Sex** | | | | | ***** |
|  | Female | 65.2% | 45.7% | 52.4% |  |
| **Ethnicity** | | | | | ***** |
|  | Hispanic | 14.1% | <20% | 42.9% |  |
| **Race** | | | | | ***** |
|  | White | 89% | 79% | 81% |  |
|  | Black OR Asian OR American Indian/ Alaska Native OR Native Hawaiian or Other Pacific Islander | <10% | 9% | <10% |  |
|  | More than one race OR other (unknown or not reported) | <10% | 12% | <10% |  |
| **Parental Education** | | | | | ***** |
|  | <HS OR High School Diploma/GED | 4% | 12% | 24% |  |
|  | Some College | 19.8% | 36.3% | 29% |  |
|  | Bachelor’s | 29.9% | 16.3% | 19% |  |
|  | Graduate | 47% | 36.3% | 29% |  |
| **Hair Toxicology Positives** | | | | |  |
|  | Nicotine Positives | 2% | <15% | <15% | * |
|  | Alcohol Positives | <1% | <1% | 0% |  |
|  | Other Substance Positives | 3% | <10% | <10% |  |
| **CBCL** |  |  |  |  |  |
|  | Internalizing Symptoms | M=49 | M=50 | M=54 | * |
|  | Externalizing Symptoms | M=44 | M=50 | M=54 | * |

Notes: Consistent with the ABCD Data User Agreement, some cells have been collapsed to obscure n<10 cells

**Table S5. Primary Neurocognitive Models Full Output**

| **Table 5a. Working Memory** | **List Sorting Working Memory Task (NIH Toolbox): Uncorrected Standard Score** | | |
| --- | --- | --- | --- |
| *Predictors* | *Estimates* | *CI* | *FDR-p* |
| (Intercept) | 75.69 | 74.07 – 77.31 | **<0.001** |
| CU Group Status | 20.50 | 12.96 – 28.04 | **<0.001** |
| Age | 1.95 | 1.89 – 2.02 | **<0.001** |
| Inverse Propensity Score | 0.79 | 0.50 – 1.08 | **<0.001** |
| Lifetime Nicotine Use | -1.21 | -1.98 – -0.45 | **0.003** |
| Lifetime Alcohol Use | 0.19 | -0.51 – 0.89 | 0.595 |
| Lifetime Other Drug Use | -0.80 | -1.51 – -0.09 | **0.04** |
| Participant's sex: Female | -0.33 | -0.87 – 0.21 | 0.26 |
| CU Group*Age | -1.32 | -1.80 – -0.84 | **<0.001** |
| **Random Effects** | | | |
| σ^2^ | 70.90 | | |
| τ_00_ _Participant ID_ | 19.94 | | |
| τ_00_ _Family ID_ | 36.64 | | |
| τ_00_ _Study Site_ | 2.98 | | |
| ICC | 0.46 | | |
| N _Participant ID_ | 4371 | | |
| N _Study Site_ | 22 | | |
| N _Family ID_ | 3668 | | |
| Observations | 13122 | | |
| Marginal R^2^ / Conditional R^2^ | 0.162 / 0.545 | | |

| **Table 5b. Inhibitory Control** | | **Flanker Inhibitory Control and Attention Task (NIH Toolbox): Uncorrected Standard Score** | | |
| --- | --- | --- | --- | --- |
| ***Predictors*** | | ***Estimates*** | ***CI*** | ***FDR-p*** |
| (Intercept) | | 72.28 | 71.40 – 73.16 | **<0.001** |
| CU Group Status | | 6.81 | 3.61 – 10.01 | **<0.001** |
| Age | | 2.12 | 2.09 – 2.16 | **<0.001** |
| Inverse Propensity Score | | 0.41 | 0.28 – 0.54 | **<0.001** |
| Lifetime Nicotine Use | | -0.72 | -1.15 – -0.29 | **0.001** |
| Lifetime Alcohol Use | | 0.10 | -0.32 – 0.52 | 0.739 |
| Lifetime Other Drug Use | | -0.06 | -0.45 – 0.33 | 0.769 |
| Participant's sex: Female | | -0.59 | -0.84 – -0.34 | **<0.001** |
| CU Group*Age | | -0.52 | -0.73 – -0.31 | **<0.001** |
| **Random Effects** | | | | |
| σ^2^ | | 38.26 | | |
| τ_00_ _Participant ID_ | | 12.53 | | |
| τ_00_ _Family ID_ | | 13.58 | | |
| τ_00_ _Study Site_ | | 1.73 | | |
| ICC | | 0.42 | | |
| N _Participant ID_ | | 10113 | | |
| N _Study Site_ | | 22 | | |
| N _Family ID_ | | 8353 | | |
| Observations | | 30039 | | |
| Marginal R^2^ / Conditional R^2^ | | 0.256 / 0.569 | | |

| **Table 5c. Episodic Memory** | | **Picture Sequence Memory Task (NIH Toolbox): Uncorrected Standard Score** | | |
| --- | --- | --- | --- | --- |
| ***Predictors*** | | ***Estimates*** | ***CI*** | ***FDR-p*** |
| (Intercept) | | 81.54 | 80.27 – 82.81 | **<0.001** |
| CU Group Status | | 16.30 | 11.24 – 21.37 | **<0.001** |
| Age | | 1.73 | 1.67 – 1.79 | **<0.001** |
| Inverse Propensity Score | | 1.07 | 0.87 – 1.27 | **<0.001** |
| Lifetime Nicotine Use | | -2.14 | -2.83 – -1.46 | **<0.001** |
| Lifetime Alcohol Use | | 0.20 | -0.47 – 0.88 | 0.557 |
| Lifetime Other Drug Use | | -0.76 | -1.38 – -0.14 | **0.024** |
| Participant's sex: Female | | 1.68 | 1.30 – 2.06 | **<0.001** |
| CU Group*Age | | -1.20 | -1.54 – -0.86 | **<0.001** |
| **Random Effects** | | | | |
| σ^2^ | | 110.52 | | |
| τ_00_ _Participant ID_ | | 27.29 | | |
| τ_00_ _Family ID_ | | 39.36 | | |
| τ_00_ _Study Site_ | | 2.61 | | |
| ICC | | 0.39 | | |
| N _Participant ID_ | | 11021 | | |
| N _Study Site_ | | 22 | | |
| N _Family ID_ | | 9113 | | |
| Observations | | 35413 | | |
| Marginal R^2^ / Conditional R^2^ | | 0.078 / 0.433 | | |

| **Table 5d. Receptive Language** | **Picture Vocabulary Task (NIH Toolbox): Uncorrected Standard Score** | | |
| --- | --- | --- | --- |
| ***Predictors*** | ***Estimates*** | ***CI*** | ***FDR-p*** |
| (Intercept) | 60.41 | 59.34 – 61.47 | **<0.001** |
| CU Group Status | 4.72 | 2.45 – 6.98 | **<0.001** |
| Age | 2.11 | 2.09 – 2.14 | **<0.001** |
| Inverse Propensity Score | 0.90 | 0.76 – 1.04 | **<0.001** |
| Lifetime Nicotine Use | -0.88 | -1.19 – -0.57 | **<0.001** |
| Lifetime Alcohol Use | -0.09 | -0.39 – 0.22 | 0.572 |
| Lifetime Other Drug Use | -0.36 | -0.65 – -0.07 | **0.017** |
| Participant's sex: Female | -0.46 | -0.72 – -0.20 | **<0.001** |
| CU Group*Age | -0.36 | -0.51 – -0.21 | **<0.001** |
| **Random Effects** | | | |
| σ^2^ | 19.45 | | |
| τ_00_ _Participant ID_ | 10.26 | | |
| τ_00_ _Family ID_ | 35.62 | | |
| τ_00_ _Study Site_ | 4.02 | | |
| ICC | 0.72 | | |
| N _Participant ID_ | 11036 | | |
| N _Study Site_ | 22 | | |
| N _Family ID_ | 9120 | | |
| Observations | 35479 | | |
| Marginal R^2^ / Conditional R^2^ | 0.241 / 0.787 | | |

| **Table 5e. Processing Speed** | **Pattern Comparison Processing Speed Task (NIH Toolbox): Uncorrected Standard Score** | | |
| --- | --- | --- | --- |
| ***Predictors*** | ***Estimates*** | ***CI*** | ***p*** |
| (Intercept) | 26.51 | 24.92 – 28.09 | **<0.001** |
| CU Group Status | 15.26 | 9.23 – 21.28 | **<0.001** |
| Age | 5.93 | 5.87 – 6.00 | **<0.001** |
| Inverse Propensity Score | 0.65 | 0.40 – 0.90 | **<0.001** |
| Lifetime Nicotine Use | -1.48 | -2.30 – -0.67 | **<0.001** |
| Lifetime Alcohol Use | 0.70 | -0.10 – 1.50 | 0.085 |
| Lifetime Other Drug Use | -1.07 | -1.80 – -0.33 | **0.005** |
| Participant's sex: Female | 2.43 | 1.96 – 2.91 | **<0.001** |
| CU Group*Age | -1.13 | -1.53 – -0.74 | **<0.001** |
| **Random Effects** | | | |
| σ^2^ | 133.36 | | |
| τ_00_ _Participant ID_ | 51.24 | | |
| τ_00_ _Family ID_ | 46.05 | | |
| τ_00_ _Study Site_ | 4.47 | | |
| ICC | 0.43 | | |
| N _Participant ID_ | 10091 | | |
| N _Study Site_ | 22 | | |
| N _Family ID_ | 8336 | | |
| Observations | 29922 | | |
| Marginal R^2^ / Conditional R^2^ | 0.434 / 0.679 | | |

| **Table 5f. Oral Reading** | **Oral Reading Recognition Task (NIH Toolbox): Uncorrected Standard Score** | | |
| --- | --- | --- | --- |
| ***Predictors*** | ***Estimates*** | ***CI*** | ***p*** |
| (Intercept) | 69.66 | 68.96 – 70.35 | **<0.001** |
| CU Group Status | 3.32 | 1.42 – 5.23 | **0.001** |
| Age | 1.89 | 1.86 – 1.91 | **<0.001** |
| Inverse Propensity Score | 0.70 | 0.58 – 0.81 | **<0.001** |
| Lifetime Nicotine Use | -0.75 | -1.01 – -0.49 | **<0.001** |
| Lifetime Alcohol Use | -0.18 | -0.43 – 0.08 | 0.228 |
| Lifetime Other Drug Use | -0.02 | -0.26 – 0.23 | 0.881 |
| Participant's sex: Female | 0.02 | -0.21 – 0.24 | 0.881 |
| CU Group*Age | -0.25 | -0.37 – -0.12 | **<0.001** |
| **Random Effects** | | | |
| σ^2^ | 13.62 | | |
| τ_00_ _Participant ID_ | 11.30 | | |
| τ_00_ _Family ID_ | 21.66 | | |
| τ_00_ _Study Site_ | 0.94 | | |
| ICC | 0.71 | | |
| N _Participant ID_ | 11020 | | |
| N _Study Site_ | 22 | | |
| N _Family ID_ | 9107 | | |
| Observations | 35339 | | |
| Marginal R^2^ / Conditional R^2^ | 0.269 / 0.790 | | |

| **Table 5g. Visuospatial** | **Little Man Task (LMT): Number correct** | | |
| --- | --- | --- | --- |
| ***Predictors*** | ***Estimates*** | ***CI*** | ***FDR-p*** |
| (Intercept) | 2.44 | 1.73 – 3.14 | **<0.001** |
| CU Group Status | 4.82 | 1.53 – 8.10 | **0.009** |
| Age | 1.53 | 1.50 – 1.56 | **<0.001** |
| Inverse Propensity Score | 0.47 | 0.37 – 0.56 | **<0.001** |
| Lifetime Nicotine Use | -0.80 | -1.15 – -0.44 | **<0.001** |
| Lifetime Alcohol Use | -0.13 | -0.51 – 0.26 | 0.631 |
| Lifetime Other Drug Use | -0.18 | -0.49 – 0.13 | 0.392 |
| Participant's sex: Female | -0.78 | -0.96 – -0.60 | **<0.001** |
| University of California, San Diego | -0.08 | -0.76 – 0.60 | 0.871 |
| University of Florida | -0.57 | -1.22 – 0.07 | 0.159 |
| University of Maryland, Baltimore | -0.11 | -0.74 – 0.51 | 0.806 |
| University of Michigan | -0.23 | -0.94 – 0.48 | 0.631 |
| University of Minnesota | 1.00 | 0.35 – 1.64 | **0.007** |
| University of Pittsburgh Medical Center | 0.34 | -0.41 – 1.09 | 0.514 |
| University of Utah | 1.56 | 0.83 – 2.30 | **<0.001** |
| University of Vermont | 0.56 | -0.13 – 1.25 | 0.206 |
| University of Wisconsin, Milwaukee | 0.34 | -0.28 – 0.97 | 0.403 |
| Virginia Commonwealth University | -0.19 | -0.89 – 0.51 | 0.685 |
| University of Colorado Boulder | -0.41 | -1.05 – 0.24 | 0.353 |
| Washington University in St. Louis | -0.03 | -0.66 – 0.60 | 0.930 |
| Yale University | 1.29 | 0.62 – 1.97 | **<0.001** |
| Icahn School of Medicine at Mount Sinai | -1.99 | -2.69 – -1.29 | **<0.001** |
| Florida International University | 1.00 | 0.40 – 1.60 | **0.003** |
| Laureate Institute for Brain Research | 0.68 | 0.02 – 1.34 | 0.087 |
| Medical University of South Carolina | 1.20 | 0.50 – 1.90 | **0.003** |
| Oregon Health & Science University | -1.00 | -1.69 – -0.32 | **0.009** |
| University of Rochester | -0.47 | -1.11 – 0.17 | 0.250 |
| SRI International | 0.03 | -0.62 – 0.67 | 0.930 |
| University of California, Los Angeles | -0.79 | -2.78 – 1.19 | 0.572 |
| CU Group * Age | -0.39 | -0.63 – -0.16 | **0.003** |
| **Random Effects** | | | |
| σ^2^ | 15.86 | | |
| τ_00_ _Participant ID_ | 7.73 | | |
| τ_00_ _Family ID_ | 8.21 | | |
| τ_00_ _Study Site_ | 0.50 | | |
| ICC | 10941 | | |
| N _Participant ID_ | 9034 | | |
| Observations | 30253 | | |
| Marginal R^2^ / Conditional R^2^ | 0.216 / 0.609 | | |

| **Table 5h. Verbal Immediate Recall** | **Rey Auditory Verbal Learning Test (RAVLT): Learning trial I - total correct** | | |
| --- | --- | --- | --- |
| ***Predictors*** | ***Estimates*** | ***CI*** | ***FDR-p*** |
| (Intercept) | 3.22 | 2.99 – 3.44 | **<0.001** |
| CU Group Status | 0.98 | -0.09 – 2.06 | 0.073 |
| Age | 0.14 | 0.13 – 0.16 | **<0.001** |
| Inverse Propensity Score | 0.10 | 0.06 – 0.13 | **<0.001** |
| Lifetime Nicotine Use | -0.17 | -0.30 – -0.03 | **0.026** |
| Lifetime Alcohol Use | 0.16 | 0.04 – 0.28 | **0.002** |
| Lifetime Other Drug Use | -0.13 | -0.24 – -0.01 | **0.041** |
| Participant's sex: Female | 0.22 | 0.15 – 0.30 | **<0.001** |
| CU Group*Age | -0.08 | -0.15 – -0.01 | **0.027** |
| **Random Effects** | | | |
| σ^2^ | 2.03 | | |
| τ_00_ _Participant ID_ | 0.29 | | |
| τ_00_ _Family ID_ | 0.58 | | |
| τ_00_ _Study Site_ | 0.05 | | |
| ICC | 0.31 | | |
| N _Participant ID_ | 4487 | | |
| N _Study Site_ | 22 | | |
| N _Family ID_ | 3786 | | |
| Observations | 13461 | | |
| Marginal R^2^ / Conditional R^2^ | 0.047 / 0.344 | | |

| **Table 5i. Verbal Memory** | **Rey Auditory Verbal Learning Test (RAVLT): Long delay trial VII - total correct** | | |
| --- | --- | --- | --- |
| ***Predictors*** | ***Estimates*** | ***CI*** | ***FDR-p*** |
| (Intercept) | 6.81 | 6.40 – 7.22 | **<0.001** |
| CU Group Status | 2.14 | 0.46 – 3.83 | **0.014** |
| Age | 0.15 | 0.14 – 0.17 | **<0.001** |
| Inverse Propensity Score | 0.17 | 0.09 – 0.24 | **<0.001** |
| Lifetime Nicotine Use | -0.50 | -0.72 – -0.29 | **<0.001** |
| Lifetime Alcohol Use | 0.02 | -0.17 – 0.21 | 0.868 |
| Lifetime Other Drug Use | -0.30 | -0.49 – -0.11 | **0.003** |
| Participant's sex: Female | 0.56 | 0.42 – 0.70 | **<0.001** |
| CU Group*Age | -0.14 | -0.25 – -0.04 | **0.011** |
| **Random Effects** | | | |
| σ^2^ | 4.45 | | |
| τ_00_ _Participant ID_ | 1.70 | | |
| τ_00_ _Family ID_ | 2.55 | | |
| τ_00_ _Study Site_ | 0.17 | | |
| ICC | 0.50 | | |
| N _Participant ID_ | 4487 | | |
| N _Study Site_ | 22 | | |
| N _Family ID_ | 3786 | | |
| Observations | 13297 | | |
| Marginal R^2^ / Conditional R^2^ | 0.028 / 0.512 | | |

**Table S6. Mean Performance on Neurocognitive Tasks by Cannabis Group and Age**

| **Category** | **Controls** | **Cannabis Use Group** |
| --- | --- | --- |
| **List Sort Working Memory [Working Memory]** | **N=3,283** | **N=1,088** |
| Age 9 | 96.38 (11.62) | 112.50 (21.92) |
| Age 10 | 99.32 (11.21) | NaN (NA) |
| Age 11 | 99.89 (12.42) | 113.00 (NA) |
| Age 12 | 105.25 (11.65) | 101.18 (6.88) |
| Age 13 | 105.39 (11.95) | 104.01 (13.18) |
| Age 14 | 107.08 (11.74) | 105.68 (11.68) |
| Age 15 | 109.08 (10.95) | 107.21 (11.17) |
| Age 16 | 110.14 (10.94) | 107.45 (11.58) |
| Age 17 | 111.07 (10.46) | 108.52 (12.86) |
| **Flanker Inhibitory Control [Inhibitory Control]** | **N=8,223** | **N=1,890** |
| Age 9 | 92.92 (9.07) | 88.80 (11.67) |
| Age 10 | 95.80 (8.56) | 92.82 (12.84) |
| Age 11 | 99.85 (7.58) | 97.97 (7.57) |
| Age 12 | 100.48 (7.74) | 100.35 (7.01) |
| Age 13 | 103.32 (7.70) | 101.80 (9.61) |
| Age 14 | 104.35 (7.63) | 102.75 (8.61) |
| Age 15 | 105.99 (7.45) | 105.43 (8.41) |
| Age 16 | 107.28 (7.09) | 105.74 (8.19) |
| Age 17 | 108.34 (6.44) | 107.34 (8.61) |
| **Picture Memory [Episodic Memory]** | **N=9,040** | **N=1,981** |
| Age 9 | 101.92 (11.83) | 101.18 (12.91) |
| Age 10 | 104.38 (12.25) | 101.67 (16.51) |
| Age 11 | 107.90 (12.53) | 106.52 (13.14) |
| Age 12 | 109.13 (12.84) | 106.44 (15.03) |
| Age 13 | 110.91 (14.71) | 106.69 (15.25) |
| Age 14 | 112.02 (15.13) | 109.25 (16.24) |
| Age 15 | 112.79 (14.80) | 108.40 (15.21) |
| Age 16 | 113.75 (14.62) | 109.12 (15.14) |
| Age 17 | 115.80 (13.47) | 111.15 (14.95) |
| **Picture Vocabulary [Receptive Language]** | **N=9,058** | **N=1,978** |
| Age 9 | 83.04 (7.53) | 77.91 (4.59) |
| Age 10 | 86.41 (8.31) | 81.42 (10.71) |
| Age 11 | 87.57 (8.20) | 86.05 (7.38) |
| Age 12 | 90.18 (8.42) | 88.25 (8.63) |
| Age 13 | 92.72 (8.83) | 89.56 (9.75) |
| Age 14 | 94.73 (8.96) | 91.92 (8.72) |
| Age 15 | 96.90 (8.37) | 94.49 (8.69) |
| Age 16 | 98.72 (8.92) | 96.06 (8.47) |
| Age 17 | 99.71 (8.40) | 98.59 (8.33) |
| **Pattern Comparison Processing Speed**  **[Processing Speed]** | **N=8,205** | **N=1,886** |
| Age 9 | 85.55 (13.89) | 86.10 (13.78) |
| Age 10 | 91.47 (14.53) | 99.36 (12.86) |
| Age 11 | 101.05 (14.43) | 98.62 (13.85) |
| Age 12 | 105.95 (15.19) | 104.73 (12.33) |
| Age 13 | 112.07 (16.10) | 106.50 (16.48) |
| Age 14 | 116.39 (16.55) | 113.95 (16.39) |
| Age 15 | 121.65 (17.08) | 119.50 (18.20) |
| Age 16 | 125.41 (17.13) | 122.68 (17.24) |
| Age 17 | 127.07 (16.70) | 123.70 (18.41) |
| **Oral Reading** | **N=9,043** | **N=1,977** |
| Age 9 | 89.78 (6.74) | 84.36 (6.82) |
| Age 10 | 92.40 (6.70) | 89.33 (6.62) |
| Age 11 | 93.86 (6.58) | 93.14 (6.82) |
| Age 12 | 95.99 (6.65) | 95.14 (6.10) |
| Age 13 | 98.39 (7.25) | 96.15 (6.96) |
| Age 14 | 99.95 (7.16) | 98.59 (7.00) |
| Age 15 | 101.25 (7.28) | 99.63 (7.23) |
| Age 16 | 103.21 (7.46) | 101.85 (7.18) |
| Age 17 | 103.31 (6.66) | 103.81 (7.52) |
| **LMT Correct [Visuospatial]** | **N=8,981** | **N=1,960** |
| Age 9 | 17.95 (5.14) | 17.55 (5.39) |
| Age 10 | 20.16 (5.62) | 20.80 (5.61) |
| Age 11 | 22.46 (6.00) | 21.65 (5.75) |
| Age 12 | 23.94 (5.94) | 22.33 (5.35) |
| Age 13 | 24.77 (5.95) | 22.79 (6.16) |
| Age 14 | 25.71 (5.76) | 24.02 (6.26) |
| Age 15 | 26.51 (5.39) | 25.93 (5.33) |
| Age 16 | 28.14 (5.21) | 24.25 (8.02) |
| Age 17 | -- | -- |
| **RAVLT Trial 1 Learning [Verbal Immediate Recall]** | **N=3,381** | **N=1,106** |
| Age 9 | 4.98 (1.79) | 3.00 (1.41) |
| Age 10 | 5.34 (1.79) | 4.00 (1.67) |
| Age 11 | 5.31 (1.62) | 5.33 (1.44) |
| Age 12 | 5.40 (1.58) | 5.52 (1.99) |
| Age 13 | 5.52 (1.61) | 5.00 (2.08) |
| Age 14 | 5.69 (1.54) | 5.52 (1.78) |
| Age 15 | 5.97 (1.75) | 5.51 (1.72) |
| Age 16 | 6.11 (1.81) | 5.53 (1.81) |
| Age 17 | 6.21 (1.66) | 6.07 (1.91) |
| **RAVLT Long Delay Memory [Verbal Memory]** | **N=3,381** | **N=1,106** |
| Age 9 | 9.15 (3.14) | 8.00 (1.41) |
| Age 10 | 9.66 (3.09) | 10.67 (3.44) |
| Age 11 | 9.33 (2.83) | 9.57 (2.96) |
| Age 12 | 9.44 (2.84) | 9.83 (2.54) |
| Age 13 | 9.09 (2.84) | 10.67 (1.21) |
| Age 14 | 10.06 (3.03) | 9.52 (3.06) |
| Age 15 | 10.23 (2.96) | 9.27 (3.13) |
| Age 16 | 10.40 (3.01) | 9.32 (3.13) |
| Age 17 | 10.77 (3.14) | 9.98 (2.79) |

**Table S7. Secondary Neurocognition-Hair Group Models Full Output**

| **Table S7a. Episodic Memory** | **Picture Sequence Memory Task (NIH Toolbox): Uncorrected Standard Score** | | |
| --- | --- | --- | --- |
| ***Predictors*** | ***Estimates*** | ***CI*** | ***p*** |
| (Intercept) | 99.56 | 84.56 – 114.56 | **<0.001** |
| Hair Cannabinoid Group [CBD] | -39.74 | -107.92 – 28.45 | 0.253 |
| Hair Cannabinoid Group [THC] | 32.69 | 6.88 – 58.50 | 0.013 |
| Age | 0.70 | 0.41 – 0.98 | **<0.001** |
| Participant's sex: Female | 3.00 | 1.26 – 4.73 | **0.001** |
| Other Positive Hair Results | -2.51 | -4.91 – -0.12 | 0.040 |
| University of Colorado Boulder | 1.13 | -13.63 – 15.89 | 0.880 |
| Florida International University | -0.78 | -15.72 – 14.17 | 0.919 |
| Laureate Institute for Brain Research | 0.84 | -13.91 – 15.58 | 0.912 |
| Medical University of South Carolina | 0.39 | -14.43 – 15.21 | 0.959 |
| Oregon Health & Science University | 3.95 | -11.44 – 19.34 | 0.615 |
| University of Rochester | -0.93 | -16.31 – 14.46 | 0.906 |
| SRI International | 7.31 | -9.52 – 24.14 | 0.394 |
| University of California, Los Angeles | -0.45 | -16.15 – 15.25 | 0.955 |
| University of California, San Diego | -2.19 | -18.01 – 13.62 | 0.786 |
| University of Florida | 0.89 | -14.13 – 15.91 | 0.908 |
| University of Maryland, Baltimore | 2.82 | -12.13 – 17.77 | 0.711 |
| University of Michigan | 0.24 | -14.65 – 15.12 | 0.975 |
| University of Minnesota | 1.74 | -13.41 – 16.90 | 0.822 |
| University of Pittsburgh Medical Center | -0.26 | -18.93 – 18.42 | 0.978 |
| University of Utah | 0.52 | -14.24 – 15.29 | 0.944 |
| University of Vermont | 4.12 | -10.67 – 18.91 | 0.585 |
| University of Wisconsin, Milwaukee | 1.49 | -13.19 – 16.16 | 0.843 |
| Virginia Commonwealth Univ | 4.20 | -12.93 – 21.33 | 0.631 |
| Washington University in St. Louis | 1.05 | -13.75 – 15.85 | 0.889 |
| Yale University | 2.89 | -12.44 – 18.22 | 0.712 |
| Hair Cannabinoid Group [CBD]*Age | 3.07 | -1.70 – 7.85 | 0.207 |
| Hair Cannabinoid Group [THC]*Age | -2.37 | -4.08 – -0.66 | **0.007** |
| **Random Effects** | | | |
| σ^2^ | 110.23 | | |
| τ_00_ _Participant ID_ | 46.15 | | |
| τ_00_ _Family ID_ | 25.31 | | |
| ICC | 0.39 | | |
| N _Participant ID_ | 645 | | |
| N _Family ID_ | 614 | | |
| Observations | 1885 | | |
| Marginal R^2^ / Conditional R^2^ | 0.040 / 0.417 | | |

| **Table S7b. Inhibitory Control** | **Flanker Inhibitory Control and Attention Task (NIH Toolbox): Uncorrected Standard Score** | | |
| --- | --- | --- | --- |
| ***Predictors*** | ***Estimates*** | ***CI*** | ***p*** |
| (Intercept) | 83.27 | 80.95 – 85.59 | **<0.001** |
| Hair Cannabinoid Group [CBD] | -29.83 | -63.71 – 4.04 | 0.084 |
| Hair Cannabinoid Group [THC] | 2.30 | -10.77 – 15.36 | 0.730 |
| Age | 1.49 | 1.34 – 1.63 | **<0.001** |
| Participant's sex: Female | -0.37 | -1.28 – 0.55 | 0.433 |
| Other Positive Hair Results | -0.90 | -2.10 – 0.31 | 0.145 |
| Hair Cannabinoid Group [CBD]*Age | 2.37 | -0.00 – 4.74 | 0.050 |
| Hair Cannabinoid Group [THC]*Age | -0.13 | -1.00 – 0.73 | 0.760 |
| **Random Effects** | | | |
| σ^2^ | 26.49 | | |
| τ_00_ _Participant ID_ | 19.67 | | |
| τ_00_ _Family ID_ | 1.89 | | |
| τ_00_ _Study Site_ | 4.52 | | |
| ICC | 0.50 | | |
| N _Study Site_ | 21 | | |
| N _Participant_ ID | 643 | | |
| N _Family ID_ | 612 | | |
| Observations | 1874 | | |
| Marginal R^2^ / Conditional R^2^ | 0.124 / 0.559 | | |

| **Table S7c. Receptive Language** | **Picture Vocabulary Task (NIH Toolbox): Uncorrected Standard Score** | | |
| --- | --- | --- | --- |
| ***Predictors*** | ***Estimates*** | ***CI*** | ***p*** |
| (Intercept) | 78.73 | 68.65 – 88.81 | **<0.001** |
| Hair Cannabinoid Group [CBD] | 13.70 | -14.15 – 41.55 | 0.335 |
| Hair Cannabinoid Group [THC] | 4.41 | -6.11 – 14.93 | 0.411 |
| Age | 1.97 | 1.86 – 2.08 | **<0.001** |
| Participant's sex: Female | -1.23 | -2.40 – -0.07 | 0.038 |
| Other Positive Hair Results | -0.98 | -2.00 – 0.04 | 0.059 |
| University of Colorado Boulder | -10.18 | -20.34 – -0.02 | 0.050 |
| Florida International University | -14.53 | -24.81 – -4.24 | 0.006 |
| Laureate Institute for Brain Research | -11.07 | -21.23 – -0.91 | 0.033 |
| Medical University of South Carolina | -8.60 | -18.81 – 1.61 | 0.099 |
| Oregon Health & Science University | -8.72 | -19.30 – 1.87 | 0.106 |
| University of Rochester | -14.59 | -25.16 – -4.01 | **0.007** |
| SRI International | -8.39 | -19.93 – 3.14 | **0.154** |
| University of California, Los Angeles | -13.04 | -23.82 – -2.25 | **0.018** |
| University of California, San Diego | -12.93 | -23.81 – -2.04 | **0.020** |
| University of Florida | -10.99 | -21.33 – -0.64 | **0.037** |
| University of Maryland, Baltimore | -6.70 | -16.99 – 3.59 | **0.202** |
| University of Michigan | -10.25 | -20.50 – -0.00 | **0.050** |
| University of Minnesota | -10.77 | -21.22 – -0.33 | **0.043** |
| University of Pittsburgh Medical Center | -14.15 | -26.95 – -1.36 | **0.030** |
| University of Utah | -8.98 | -19.15 – 1.19 | 0.083 |
| University of Vermont | -6.45 | -16.64 – 3.74 | 0.214 |
| University of Wisconsin, Milwaukee | -7.84 | -17.95 – 2.27 | 0.128 |
| Virginia Commonwealth Univ | -10.01 | -21.77 – 1.75 | 0.095 |
| Washington University in St. Louis | -10.31 | -20.51 – -0.11 | 0.048 |
| Yale University | -6.78 | -17.33 – 3.77 | 0.208 |
| Hair Cannabinoid Group [CBD]*Age | -0.93 | -2.88 – 1.02 | 0.352 |
| Hair Cannabinoid Group [THC]*Age | -0.38 | -1.07 – 0.32 | 0.290 |
| **Random Effects** | | | |
| σ^2^ | 16.54 | | |
| τ_00_ _Participant ID_ | 10.74 | | |
| τ_00_ _Family ID_ | 33.82 | | |
| ICC | 0.73 | | |
| N _Participant ID_ | 644 | | |
| N _Family ID_ | 613 | | |
| Observations | 1877 | | |
| Marginal R^2^ / Conditional R^2^ | 0.220 / 0.789 | | |

| **Table S7d. Oral Reading** | **Oral Reading Recognition Task (NIH Toolbox): Uncorrected Standard Score** | | |
| --- | --- | --- | --- |
| ***Predictors*** | ***Estimates*** | ***CI*** | ***p*** |
| (Intercept) | 74.76 | 72.98 – 76.55 | **<0.001** |
| Hair Cannabinoid Group [CBD] | 8.91 | -17.02 – 34.83 | 0.500 |
| Hair Cannabinoid Group [THC] | 12.35 | 2.55 – 22.15 | 0.014 |
| Age | 1.81 | 1.71 – 1.92 | **<0.001** |
| Participant's sex: Female | -0.74 | -1.76 – 0.28 | 0.153 |
| Other Positive Hair Results | -1.00 | -1.95 – -0.05 | 0.040 |
| Hair Cannabinoid Group [CBD]*Age | -0.64 | -2.46 – 1.17 | 0.488 |
| Hair Cannabinoid Group [THC]*Age | -0.82 | -1.47 – -0.17 | 0.013 |
| **Random Effects** | | | |
| σ^2^ | 14.44 | | |
| τ_00_ _Participant ID_ | 11.71 | | |
| τ_00_ _Family ID_ | 22.20 | | |
| τ_00_ _Study Site_ | 1.53 | | |
| ICC | 0.71 | | |
| N _Participant ID_ | 643 | | |
| N _Study Site_ | 21 | | |
| N _Family ID_ | 612 | | |
| Observations | 1871 | | |
| Marginal R^2^ / Conditional R^2^ | 0.172 / 0.760 | | |

| **Table S7e. Processing Speed** | **Pattern Comparison Processing Speed Task (NIH Toolbox): Uncorrected Standard Score** | | |
| --- | --- | --- | --- |
| ***Predictors*** | ***Estimates*** | ***CI*** | ***p*** |
| (Intercept) | 24.95 | 7.74 – 42.16 | **0.005** |
| Hair Cannabinoid Group [CBD] | -45.13 | -115.46 – 25.20 | 0.208 |
| Hair Cannabinoid Group [THC] | -5.84 | -33.17 – 21.50 | 0.675 |
| Age | 5.16 | 4.87 – 5.45 | <0.001 |
| Participant's sex: Female | 1.69 | -0.32 – 3.71 | 0.100 |
| Other Positive Hair Results | -3.62 | -6.13 – -1.10 | **0.005** |
| University of Colorado Boulder | 19.66 | 2.59 – 36.73 | 0.024 |
| Florida International University | 15.54 | -1.73 – 32.82 | 0.078 |
| Laureate Institute for Brain Research | 15.87 | -1.19 – 32.93 | 0.068 |
| Medical University of South Carolina | 16.96 | -0.18 – 34.11 | 0.052 |
| Oregon Health & Science University | 20.60 | 2.81 – 38.39 | 0.023 |
| University of Rochester | 15.81 | -1.98 – 33.61 | 0.082 |
| SRI International | 30.00 | 10.59 – 49.41 | **0.002** |
| University of California, Los Angeles | 5.58 | -12.57 – 23.73 | 0.547 |
| University of California, San Diego | 9.26 | -9.08 – 27.59 | 0.322 |
| University of Florida | 13.19 | -4.20 – 30.57 | 0.137 |
| University of Maryland, Baltimore | 14.88 | -2.41 – 32.17 | 0.092 |
| University of Michigan | 21.01 | 3.79 – 38.22 | 0.017 |
| University of Minnesota | 17.70 | 0.16 – 35.24 | 0.048 |
| University of Pittsburgh Medical Center | 9.44 | -12.16 – 31.03 | 0.391 |
| University of Utah | 15.98 | -1.10 – 33.07 | 0.067 |
| University of Vermont | 20.67 | 3.56 – 37.79 | 0.018 |
| University of Wisconsin, Milwaukee | 20.41 | 3.44 – 37.38 | 0.018 |
| Virginia Commonwealth Univ | 14.97 | -4.91 – 34.84 | 0.140 |
| Washington University in St. Louis | 19.07 | 1.95 – 36.18 | 0.029 |
| Yale University | 20.83 | 3.10 – 38.55 | 0.021 |
| Hair Cannabinoid Group [CBD]*Age | 3.59 | -1.33 – 8.52 | 0.152 |
| Hair Cannabinoid Group [THC]*Age | 0.23 | -1.58 – 2.04 | 0.802 |
| **Random Effects** | | | |
| σ^2^ | 112.23 | | |
| τ_00_ _Participant ID_ | 55.05 | | |
| τ_00_ _Family ID_ | 52.36 | | |
| ICC | 0.49 | | |
| N _Participant ID_ | 642 | | |
| N _Family ID_ | 611 | | |
| Observations | 1863 | | |
| Marginal R^2^ / Conditional R^2^ | 0.310 / 0.647 | | |

**Table S8. Mean Performance on Neurocognitive Tasks by Hair Cannabinoid Group and Age**

| **Category** | **Controls** | **THC Only Group** | **CBD+ Group** |
| --- | --- | --- | --- |
| **Picture Memory [Episodic Memory]** | | |  |
| Age 10 | 111.11 (15.34) |  |  |
| Age 11 | 110.58 (11.60) | 114.00 (12.06) |  |
| Age 12 | 110.65 (11.27) | 111.83 (20.84) | 106.50 (3.54) |
| Age 13 | 112.65 (14.08) | 108.17 (12.77) | 99.00 (15.24) |
| Age 14 | 113.89 (14.54) | 110.55 (12.66) | 116.40 (13.69) |
| Age 15 | 113.78 (14.15) | 107.56 (14.57) | 116.00 (4.00) |
| Age 16 | 113.50 (14.56) | 104.54 (13.99) |  |
| Age 17 | 113.57 (17.09) | 102.33 (16.50) | 106.00 |
| **Pattern Comparison Processing Speed [Processing Speed]** | | |  |
| Age 10 | 98.04 (12.51) |  |  |
| Age 11 | 102.49 (13.10) | 88.25 (11.32) |  |
| Age 12 | 107.73 (14.60) | 102.00 (8.25) | 108.00 (19.80) |
| Age 13 | 115.97 (14.46) | 113.64 (8.91) | 114.43 (19.97) |
| Age 14 | 119.64 (14.74) | 114.05 (18.53) | 118.00 (6.63) |
| Age 15 | 123.67 (16.04) | 121.94 (17.65) | 119.00 (14.18) |
| Age 16 | 126.97 (16.00) | 119.31 (20.67) |  |
| Age 17 | 126.71 (21.45) | 132.50 (17.68) | 155.00 |
| **Picture Vocabulary [Receptive Language]** | | |  |
| Age 10 | 92.43 (8.28) |  |  |
| Age 11 | 89.89 (7.50) | 86.25 (5.74) |  |
| Age 12 | 93.29 (8.05) | 91.00 (5.51) | 97.50 (2.12) |
| Age 13 | 95.67 (8.54) | 96.00 (8.18) | 95.83 (10.70) |
| Age 14 | 97.49 (7.96) | 95.65 (8.80) | 92.00 (8.52) |
| Age 15 | 99.07 (7.85) | 93.50 (6.46) | 102.00 (4.36) |
| Age 16 | 100.47 (8.19) | 94.46 (8.75) |  |
| Age 17 | 99.75 (7.03) | 104.33 (7.37) | 87.00 |
| **Flanker Inhibitory Control [Inhibitory Control]** | | | |
| Age 10 | 100.75 (7.03) |  |  |
| Age 11 | 100.27 (7.19) | 99.25 (6.95) |  |
| Age 12 | 101.93 (6.81) | 101.60 (5.41) | 95.00 (4.24) |
| Age 13 | 104.63 (6.98) | 107.55 (4.48) | 106.29 (6.99) |
| Age 14 | 105.61 (6.37) | 105.16 (6.23) | 104.89 (3.52) |
| Age 15 | 106.93 (6.90) | 107.47 (7.77) | 110.33 (3.51) |
| Age 16 | 107.43 (7.58) | 104.92 (8.28) |  |
| Age 17 | 107.26 (12.99) | 112.33 (3.79) | 110.00 |
| **Oral Reading** | | | |
| Age 10 | 95.61 (6.00) |  |  |
| Age 11 | 94.74 (6.44) | 94.00 (8.04) |  |
| Age 12 | 97.27 (6.76) | 96.33 (4.80) | 91.00 (0.00) |
| Age 13 | 99.56 (7.56) | 102.17 (4.88) | 99.50 (4.04) |
| Age 14 | 100.75 (6.83) | 101.50 (6.25) | 97.60 (6.22) |
| Age 15 | 102.41 (7.29) | 99.34 (4.34) | 99.00 (8.19) |
| Age 16 | 104.20 (7.48) | 102.00 (9.21) |  |
| Age 17 | 104.28 (6.96) | 105.33 (3.79) | 97.00 |

1.	Cooper GA, Kronstrand R, Kintz P, Society of Hair T. Society of Hair Testing guidelines for drug testing in hair. *Forensic Sci Int.* 2012;218(1-3):20-24.

2.	Hill VA, Schaffer MI, Stowe GN. Carboxy-THC in Washed Hair: Still the Reliable Indicator of Marijuana Ingestion. *J Anal Toxicol.* 2016;40(5):345-349.

3.	van Buuren S, Groothuis-Oudshoorn K. mice: Multivariate Imputation by Chained Equations in R. *Journal of Statistical Software.* 2011;45:1-67.
